# Co-evolution of Oncogenic KRAS Signaling and LILRB^high^ Macrophages Drives Pancreatic Cancer Recurrence

**DOI:** 10.64898/2026.03.05.709991

**Authors:** Jianzhen Lin, Zhenzhen Xun, Fangze Qian, Zhigao Chen, Weikang Hu, Wei Liu, Yang Wu, Hao Yuan, Lingdi Yin, Yazhou Wang, Xiao Huang, Yujia Dang, Bin Xiao, Junli Wu, Wentao Gao, Jishu Wei, Qiang Li, Min Tu, Jing Zhou, Xu Feng, Zipeng Lu, Li Wen, Kuirong Jiang, Han Liang

**Affiliations:** Pancreas Center, The First Affiliated Hospital of Nanjing Medical University & Jiangsu Province Hospital; Pancreas Institute, Nanjing Medical University, Nanjing, China; Department of Bioinformatics and Computational Biology, The University of Texas MD Anderson Cancer Center, Houston, Texas, USA; Center for Biomarker Discovery and Validation, Institute of Clinical Medicine, Peking Union Medical College Hospital, Chinese Academy of Medical Science, Beijing, China; Nanjing Leads Biolabs Co., Ltd., Nanjing, Jiangsu Province, China; Jiangsu Provincial Key Laboratory of Chronic Digestive Diseases, The First Affiliated Hospital of Nanjing Medical University, Nanjing, China; Department of Systems Biology, The University of Texas MD Anderson Cancer Center, Houston, Texas, USA

## Abstract

Pancreatic ductal adenocarcinoma (PDAC) frequently recurs after surgical resection, indicating that residual disease is sustained by coordinated tumor–microenvironment interactions. To define the biological basis of recurrence, we leverage large-scale clinical data from 2,710 patients, deeply characterized multi-omics profiling (whole-exome, bulk RNA, and single-nucleus sequencing) of 36 matched primary and locally recurrent PDACs, an in-house multiplex spatial imaging cohort of 190 patients, and extensive public datasets. Recurrent tumors were characterized by increased *KRAS* mutant allele dosage and reinforced KRAS signaling, accompanied by expansion of basal-like malignant cell states. In parallel, we identified an immunosuppressive macrophage population marked by high LILRB expression that spatially co-localized with KRAS-activated tumor cells. Functional studies showed that LILRB4^+^ macrophages enhanced tumor cell plasticity and progression, whereas inhibition of macrophage LILRB4 suppressed these phenotypes. Notably, a first-in-class human anti-LILRB4 antibody reduced macrophage-driven tumor traits, and dual targeting of KRAS signaling and LILRB4 achieved superior tumor control in macrophage-containing mouse models. These findings reveal a co-evolved tumor–immune niche underlying PDAC recurrence and nominate the KRAS–LILRB4 axis as a therapeutic vulnerability.

## Introduction

Pancreatic ductal adenocarcinoma (PDAC) remains one of the deadliest malignancies, with a 5-year survival rate of approximately 13% and a projected rise to the second leading cause of cancer-related mortality within the next decade^1^. Surgical resection offers the only potential cure, yet recurrence is common: 20–30% of patients relapse within two years, and up to 80% experience recurrence within five years. These observations indicate that residual disease persists despite apparently curative surgery and underscore the need to define the biological mechanisms that sustain postoperative relapse.

Genomic studies have established PDAC evolution as a hybrid process involving both stepwise and punctuated events, with *KRAS* mutations serving as a central oncogenic driver^2,3^. Detailed clonal analyses have delineated trajectories of metastatic spread and informed emerging immunotherapeutic approaches, including neoantigen-targeted strategies^4,5^. However, current understanding remains heavily focused on tumor-intrinsic alterations. The consequences of *KRAS* mutant allele dosage in recurrent disease remain insufficiently explored, and the dynamic remodeling of the tumor microenvironment (TME) during recurrence is poorly defined.

PDAC is characterized by a dense and immunosuppressive microenvironment dominated by myeloid populations that can constrain antitumor immunity and promote tumor progression^6^. Yet most available data derive from cross-sectional analyses of primary or metastatic lesions sampled from distinct anatomical sites. Such approaches are limited in their ability to capture the spatiotemporal evolution of the immune ecosystem during relapse, particularly in the context of residual oncogenic signaling. Consequently, how tumor-intrinsic signaling and immune remodeling co-evolve to drive recurrence remains unclear.

Recent advances in single-cell and spatial transcriptomics have enabled high-resolution interrogation of tumor–immune interactions. A longitudinal framework comparing matched primary and locally recurrent tumors from the same patient and anatomical site offers a unique opportunity to define the cellular and molecular programs that specifically underpin postoperative relapse. Here, we performed whole-exome sequencing, bulk RNA sequencing, single-nucleus RNA-sequencing, and multiplex spatial imaging of 36 matched primary and locally recurrent PDACs. By integrating these datasets, we delineate the coordinated evolution of tumor-intrinsic KRAS signaling and the immune microenvironment during recurrence. Our findings reveal a recurrence-associated ecosystem characterized by amplified KRAS activity and expansion of immunosuppressive macrophage states, providing mechanistic insight into PDAC relapse and identifying actionable vulnerabilities for therapeutic intervention.

## Results

### Study overview and patient characteristics

To assess the clinical impact of recurrence, we assembled a cohort of 2,710 patients who underwent surgical resection for pancreatic cancer between December 2014 and December 2024 (**Tables S1–S3**). The survival divergence was striking: patients who developed recurrence had a median survival of only 577 days, whereas about 90% (89.32%) of recurrence-free patients remained alive beyond 3,000 days (log-rank, *P* < 10^-22^; **Fig. 1A**). Importantly, after adjustment for age, sex, pathological stage and receipt of adjuvant chemotherapy, recurrence remained the strongest independent predictor of poor outcome in multivariable Cox analyses (*P* < 1 x 10^-3^; **Fig. 1B** and **Fig. S1**). These findings establish recurrence as a dominant determinant of pancreatic cancer mortality and highlight the urgent need to elucidate its underlying biological drivers.

**Figure 1.**
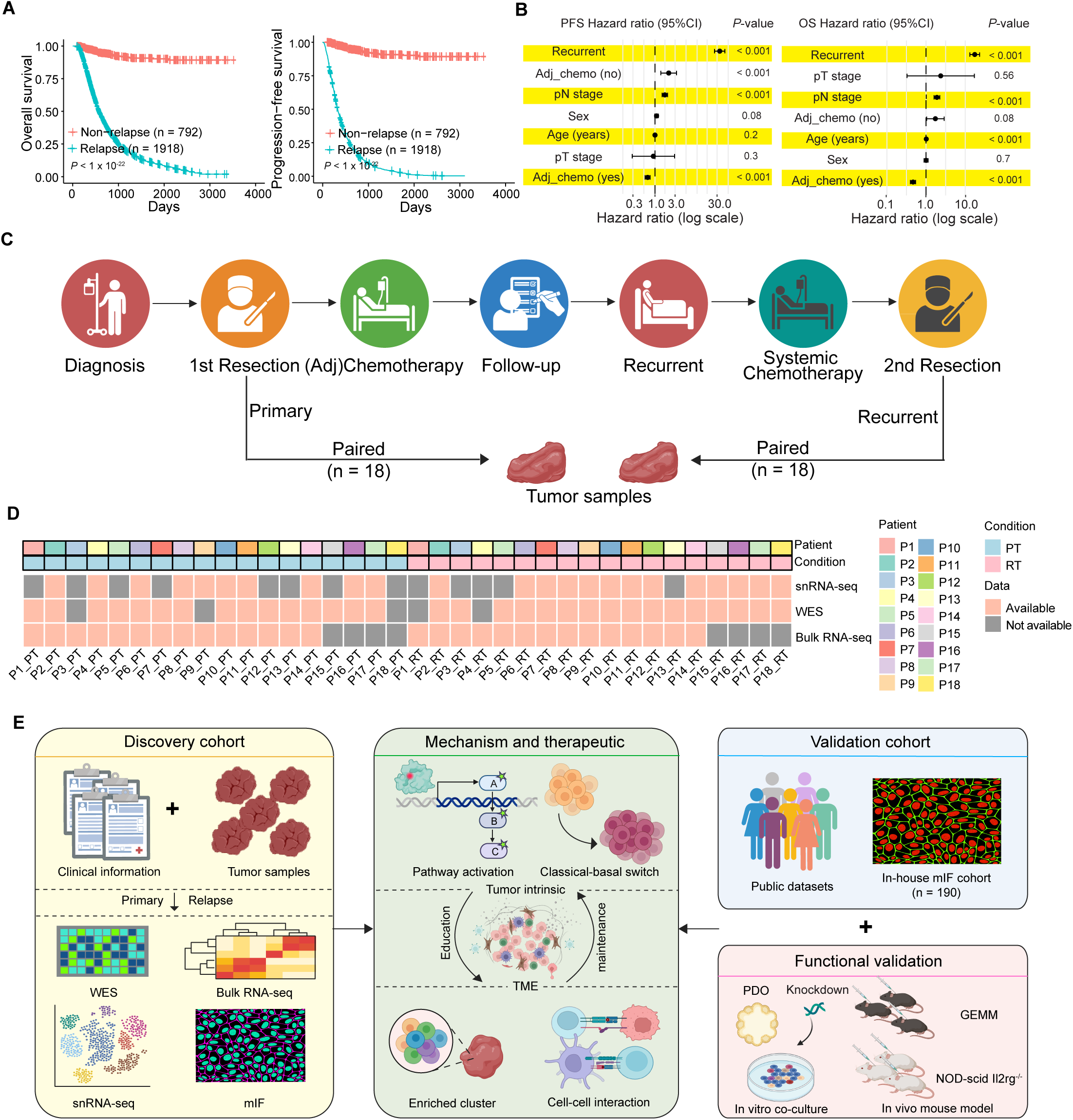
Study design and data overview. **(A)** Kaplan–Meier curves of overall survival (OS, left) and progression-free survival (PFS, right) for patients who remained recurrence-free (Non-relapse, *n* = 792) or developed recurrence (Relapse, *n* = 1,918) in a PDAC clinical cohort; *P* values, log-rank test. **(B)** Multivariable Cox regression analyses for PFS (left) and OS (right). Forest plots show hazard ratios (HRs) with 95% confidence intervals (CIs) for recurrence status and clinical covariates (age, sex, pathological T stage [pT stage], and adjuvant chemotherapy); *P* values from Cox models. **(C)** Clinical timeline and biospecimen collection scheme. Primary tumors were obtained at first resection; recurrent tumors were collected at second resection when feasible, yielding 18 matched primary–recurrent pairs. **(D)** Data availability map across the 36 paired primary-recurrent cases, indicating patient identity (P1–P18), sampling condition (primary, PT; recurrent, RT), and the presence or absence of each profiling modality (snRNA-seq, WES, bulk RNA-seq). **(E)** Overview of the integrated study framework. The discovery cohort integrated clinical annotation with multi-modal profiling (WES, bulk RNA-seq, snRNA-seq, and multiplex immunofluorescence [mIF]) to define tumor-intrinsic programs, classical–basal state switching, and remodeling of the tumor microenvironment. We then performed validation using public datasets and an independent in-house whole-slide mIF cohort (n = 190). We further carried out functional validation *in vitro* using patient-derived organoid (PDO) co-culture and gene knockdown assays and in vivo using genetically engineered mouse models (GEMMs) and NOD-scid Il2rg^−/−^ mouse models. OS, overall survival; PFS, progression-free survival; HR, hazard ratio; CI, confidence interval; WES, whole-exome sequencing; snRNA-seq, single-nucleus RNA sequencing; mIF, multiplex immunofluorescence. See also **Fig. S1** and **Tables S1-3**.

To define the biological basis of PDAC recurrence, we focused on patients with resectable disease at initial diagnosis (**Table S2**) and established a longitudinal sampling framework. Primary tumors were collected at first resection, followed by standard adjuvant therapy and structured surveillance. For patients who subsequently developed recurrence and were eligible for further surgery, recurrent tumors were obtained at second resection when clinically feasible. This paired, same-patient and same-anatomical-site design minimizes interpatient and organ-specific confounders, enabling direct interrogation of tumor and microenvironmental evolution during relapse. In total, we generated 18 matched primary–recurrent tumor pairs comprising our discovery cohort (**Fig. 1C, D**).

Leveraging these longitudinal specimens, we performed multi-omics profiling—including whole-exome sequencing, bulk RNA sequencing, single-nucleus RNA sequencing (snRNA-seq), and multiplex spatial imaging (**Fig. 1D, E**). We defined tumor-intrinsic evolution by assessing oncogenic pathway activation and malignant cell state plasticity, while concurrently resolving recurrence-associated remodeling of the TME through identification of enriched immune populations and tumor–immune interactions. Findings were further validated in independent public single-cell and spatial transcriptomic datasets and in an in-house whole-slide multiplex immunofluorescence (mIF) cohort (n = 190) (**Fig. 1E**). Candidate therapeutic vulnerabilities were functionally interrogated using a PDAC patient-derived organoid (PDO) organoid–macrophage co-culture system, an immunocompetent KPC genetically engineered mouse model (GEMM), and a subcutaneous tumor–macrophage xenograft model. Collectively, by combining longitudinal multi-omics profiling with multi-layer functional interrogation, we delineate coordinated tumor–immune evolution as a central mechanism of PDAC recurrence and uncover actionable therapeutic vulnerabilities.

### Enhanced KRAS signaling and basal reprogramming in recurrence

To define somatic alterations associated with recurrence, we analyzed whole-exome sequencing data from matched primary–recurrent tumor pairs. Among established PDAC driver genes^7^, *KRAS* was the only gene exhibiting a significant increase in variant allele frequency (VAF) in recurrent tumors compared with matched primaries (*P* = 0.05; **Fig. 2A** and **Fig. S2A**). *KRAS* mutations were also more frequent in recurrent lesions (**Fig. S2B**), and *KRAS* showed the most pronounced increase in mutant allele dosage among all examined genes (*P* = 0.03; **Fig. 2B**). In the TCGA PDAC cohort, high *KRAS* mutant allele dosage was associated with significantly shorter overall survival (log-rank, *P* = 0.04; **Fig. 2C**). Collectively, these data suggest positive selection for increased *KRAS* mutant allele dosage during relapse, indicating that reinforced oncogenic signals represent a defining feature of recurrent PDAC evolution.

**Figure 2.**
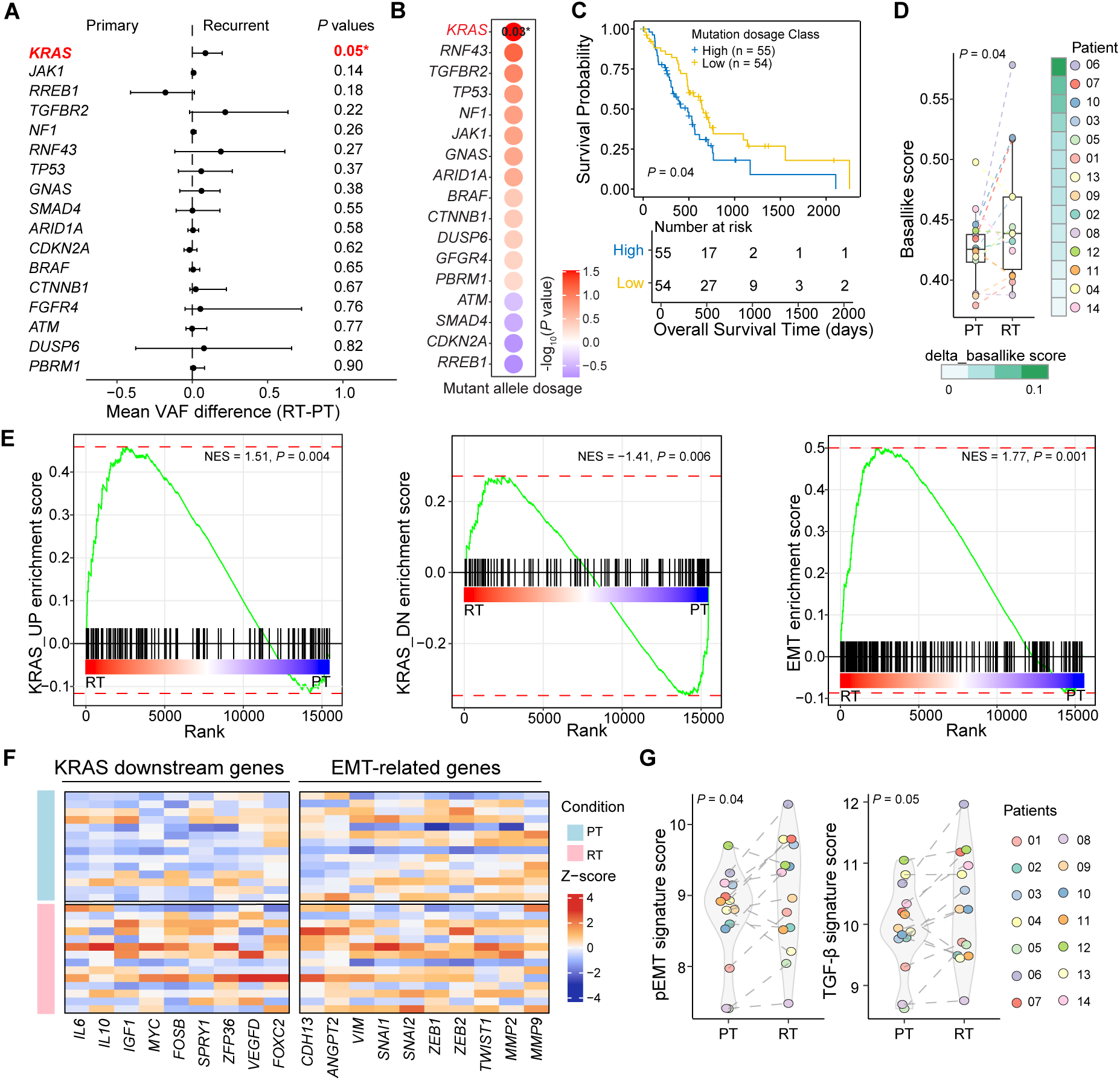
Bulk sequencing reveals increased *KRAS* mutant allele dosage, basal-like state and KRAS pathway activation in recurrent PDAC. **(A)** Gene-level changes in variant allele frequency (VAF) between primary tumors (PT) and matched recurrent tumors (RT). Points show mean VAF difference (RT–PT) across paired samples for common PDAC driver genes; error bars indicate uncertainty of the mean; *P* values from paired comparisons are shown. *P* value, t-test. **(B)** Changes in mutant allele dosage between RT and PT across paired samples. Dot color indicates signed statistical significance. *P* value, single sided t-test. **(C)** Kaplan–Meier analysis of overall survival in TCGA-PDAC stratified by KRAS mutant allele dosage (High vs Low; median split). *P* value, log-rank test. **(D)** Basal-like transcriptomic scores in paired PT and RT tumors profiled by bulk RNA-seq; dots represent individual samples; paired samples are connected. For the heatmap, the color represents the difference of basal-like score from RT minus basal-like score from PT*. P* value, single sided t-test. **(E)** Gene set enrichment analysis (GSEA) of bulk RNA-seq comparing RT versus PT, showing enrichment of KRAS up-regulation, KRAS down-regulation, and Hallmark EMT signatures; normalized enrichment score (NES) and *P* values are indicated. **(F)** Heatmaps of KRAS downstream genes (left) and EMT downstream genes (right) across PT and RT samples; values are Z-scored per gene. **(G)** Paired mean signature scores for partial EMT (pEMT) and TGF-β programs based on the bulk RNA-seq data of PT and RT samples; dots represent samples and lines connect matched pairs; *P* values from paired comparisons. PT, primary tumor; RT, recurrent tumor; VAF, variant allele frequency; NES, normalized enrichment score; EMT, epithelial–mesenchymal transition; pEMT, partial EMT. See also **Fig. S2**.

Increased *KRAS* mutant allele dosage is frequently associated with heightened KRAS pathway activation^8^ and transition from a classical to a basal-like transcriptional state^9^. We therefore asked whether recurrent tumors exhibit enhanced KRAS signaling and basal-like reprogramming. Using bulk RNA sequencing, we quantified lineage states and observed significantly higher basal-like scores in recurrent tumors compared with matched primaries (*P* < 0.04; **Fig. 2D**), a finding confirmed by categorical subtype assignment^10^ (**Fig. S2C**). This shift toward a basal-like state is consistent with more aggressive biology and poorer clinical outcomes.

Gene set enrichment analysis further demonstrated strong enrichment of KRAS-upregulated signatures in recurrent tumors, whereas KRAS-downregulated signatures were enriched in primary tumors (**Fig. 2E**). Recurrent lesions also displayed marked enrichment of Hallmark epithelial–mesenchymal transition (EMT) programs, which supports higher plasticity and a more basal-like program underlying recurrence^11–13^ (**Fig. 2E, F**), alongside increased partial EMT (pEMT) and

TGFβ signature scores (**Fig. 2G**), as well as elevated TNFα, EMT and KRAS signaling activity (**Fig. S2D**). Together, these findings indicate convergence of increased KRAS mutant allele dosage, amplified KRAS pathway activation and EMT-associated plasticity during recurrence, nominating KRAS-driven transcriptional reprogramming as a central and potentially tractable axis in recurrent PDAC.

### Expansion of a KRAS-activated basal-like state drives PDAC recurrence

To resolve tumor evolution at single-cell resolution, we analyzed snRNA-seq data from the 18 matched primary and recurrent PDAC pairs. After rigorous quality control and Harmony-based^14^ integration, 71,541 cells from primary tumors and 82,464 cells from recurrent tumors were retained. Marker-based annotation identified 11 major cell populations, including epithelial, ductal, acinar, endocrine, macrophage, T cell, B cell, dendritic cell, plasma, endothelial and fibroblast compartments (**Fig. 3A** and **Fig. S3A**). Whole-slide mIF profiling of matched specimens confirmed comparable overall cell-type distributions between primary and recurrent tumors (**Fig. 3B** and **Fig. S3B**), indicating that recurrence is not driven by gross compositional shifts.

**Figure 3.**
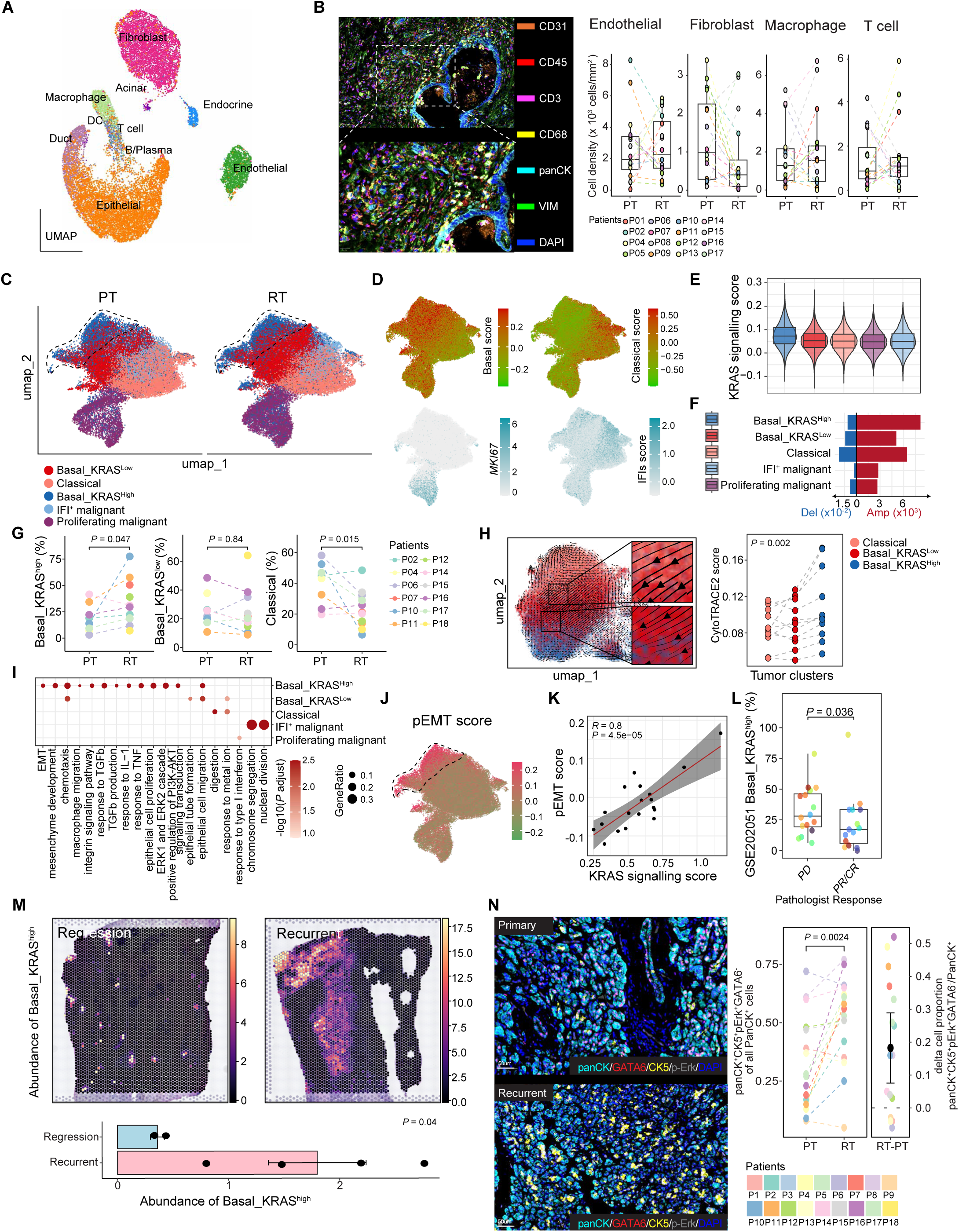
Single-nuclear RNA-sequencing analysis reveals enrichment of KRAS-activated basal malignant cells in recurrent PDAC. **(A)** UMAP projection of integrated snRNA-seq profiles from matched primary tumors (PT; 71,541 cells) and recurrent tumors (RT; 82,464 cells) across patients, annotated by major cell types. **(B)** Representative whole-slide multiplex immunofluorescence (mIF) images from matched PT and RT samples (left) stained for CD31 (orange), CD45 (red), CD3 (magenta), CD68 (yellow), PanCK (light blue), VIM (green), and DAPI (blue), with quantification of endothelial cells (CD31^+^VIM^+^), fibroblasts (VIM^+^CD41^-^CD45-PanCK^-^), macrophages (CD68^+^), and T cells (CD3^+^) as tissue-level cell density across paired samples (right). **(C)** UMAP of inferCNV-defined malignant epithelial cells, split by PT and RT, colored by malignant states (Basal_KRAS^high^, Basal_KRAS^low^, Classical, IFI^+^ malignant, and Proliferating malignant). **(D)** Feature maps of basal score, classical score, proliferation (MKI67), and interferon-stimulated gene (IFI) score across malignant cells. **(E)** KRAS signaling score across malignant states (violin plots). **(F)** KRAS copy-number variation (CNV) state scores across malignant states, including amplification and deletion components. **(G)** Paired changes in the fraction of malignant cells in Basal_KRAS^high^, Basal_KRAS^low^, and Classical states from PT to RT across patients; lines connect matched samples. *P* value, single-sided t-test. **(H)** Trajectory inference from Classical through Basal_KRAS^low^ to Basal_KRAS^high^ (left) with CytoTRACE2 scores across malignant states (right). **(I)** Dot plot of pathway enrichment across malignant states; dot size indicates gene ratio and color denotes significance. **(J)** Feature map of partial EMT (pEMT) score across malignant cells. **(K)** Association between KRAS signaling score and pEMT score across malignant cells; line indicates linear fit with 95% CI; correlation coefficient (*R*) and *P* value are shown. **(L)** External validation in GSE202051dataset: fraction of Basal_KRAS^high^ malignant cells in tumors with poor pathological response (PD) versus partial/complete response (PR/CR). **(M)** Spatial validation in HTAN-WUSTL PDAC dataset: Basal_KRAS^high^ infiltration in regression (after neoadjuvant 6*FOLFIRINOX) versus recurrent (under adjuvant chemoradiotherapy) samples (left), with quantification (right). **(N)** Representative mIF validation of Basal_KRAS^high^ (panCK^+^ CK5^+^ p-Erk^+^ GATA6^-^) infiltration in matched PT and RT samples (left) stained for p-ERK (yellow), GATA6 (red), PanCK (light blue), CK5 (dark green), and DAPI (blue); scale bars, 100 µm. Quantification of Basal_KRAS^high^ proportion of all PanCK^+^ cells in matched PT and RT samples (left) and paired change (RT–PT) (right); lines connect matched samples; *P* value from paired t-test. PT, primary tumor; RT, recurrent tumor; mIF, multiplex immunofluorescence; CNV, copy-number variation; IFIs, interferon-stimulated genes; EMT, epithelial–mesenchymal transition; pEMT, partial EMT. See also **Fig. S3**.

We next focused on malignant cell heterogeneity. After distinguishing malignant from normal epithelial cells using inferCNV^15^ (**Fig. S3C**), we defined five malignant states based on basal and classical lineage markers^16^ and KRAS pathway activity: Basal_KRAS^high^, Basal_KRAS^low^, Classical, Proliferative and IFI^+^ states (**Fig. 3C–E** and **Fig. S3D**). Basal_KRAS^high^ cells exhibited the highest *KRAS* copy-number amplification scores (**Fig. 3F**). Notably, recurrent tumors showed significant expansion of Basal_KRAS^high^ cells (*P* = 0.047) and contraction of Classical cells (*P* = 0.015) (**Fig. 3G** and **Fig. S3E**), consistent with bulk transcriptomic evidence of basal reprogramming. CytoTRACE2^17^ analysis inferred a trajectory from Classical to Basal_KRAS^low^ and ultimately to Basal_KRAS^high^ states within individual patients (**Fig. 3H**), supporting a state transition during recurrence.

Functionally, Basal_KRAS^high^ cells were enriched for KRAS signaling, EMT, TGFβ, TNFα and inflammatory pathways (**Fig. 3I** and **Fig. S3F**). These cells displayed elevated pEMT scores^18^, which strongly correlated with KRAS pathway activity across samples (R = 0.8, *P* = 4.5 × 10⁻⁵; **Fig. 3J,K**). Moreover, recurrence-associated Basal_KRAS^high^ cells upregulated immune-modulatory factors (e.g., *FN1*) (**Fig. S3G**), linking oncogenic signaling with microenvironmental crosstalk.

Independent validation across multiple cohorts confirmed these findings. In a chemotherapy-treated PDAC dataset (GSE202051^19^), poor responders exhibited higher proportions of Basal_KRAS^high^ cells than responders (*P* = 0.036; **Fig. 3L**, and **Fig. S3H-K**). Similarly, recurrent cases in the HTAN-WUSTL PDAC cohort^20^ showed increased Basal_KRAS^high^ enrichment compared with regression samples (*P* = 0.04; **Fig. 3M**). Consistent patterns were observed in our independent mIF validation cohort (*P* = 0.0024, **Fig. 3N**).

Collectively, these results identify expansion of a KRAS-activated basal-like malignant state as a defining feature of PDAC recurrence, linking oncogenic dosage amplification to transcriptional plasticity and immune-interactive programs.

### LILRB^high^ macrophages couple with KRAS-activated basal cells in recurrence

To define tumor-immune crosstalk, we performed LIANA-based^21^ ligand–receptor analysis and identified macrophages as the most interactive immune compartments with malignant cells (**Fig. 4A**). Given this observation and the established role of tumor-associated macrophages (TAMs) in shaping the PDAC microenvironment^22–24^, we next interrogated macrophage heterogeneity in recurrence. Single-cell analysis identified ten macrophage subtypes (**Fig. 4B** and **Fig. S4A**). One distinct subset exhibited high expression of LILRB family members (*LILRB1–5*) and was designated LILRB^high^ macrophages. LILRB receptors contain immunoreceptor tyrosine-based inhibitory motifs^25,26^ and are associated with immunosuppressive signaling. Consistently, LILRB^high^ cells expressed canonical pro-tumor markers, including *CD163* and *MRC1*^27–29^ (**Fig. 4C**). Among all macrophage subsets, only LILRB^high^ macrophages were significantly enriched in recurrent tumors relative to matched primaries (*P* < 0.014, **Fig. 4D** and **Fig. S4B–D**) and displayed the highest immunosuppressive signature scores (**Fig. 4E**).

**Figure 4.**
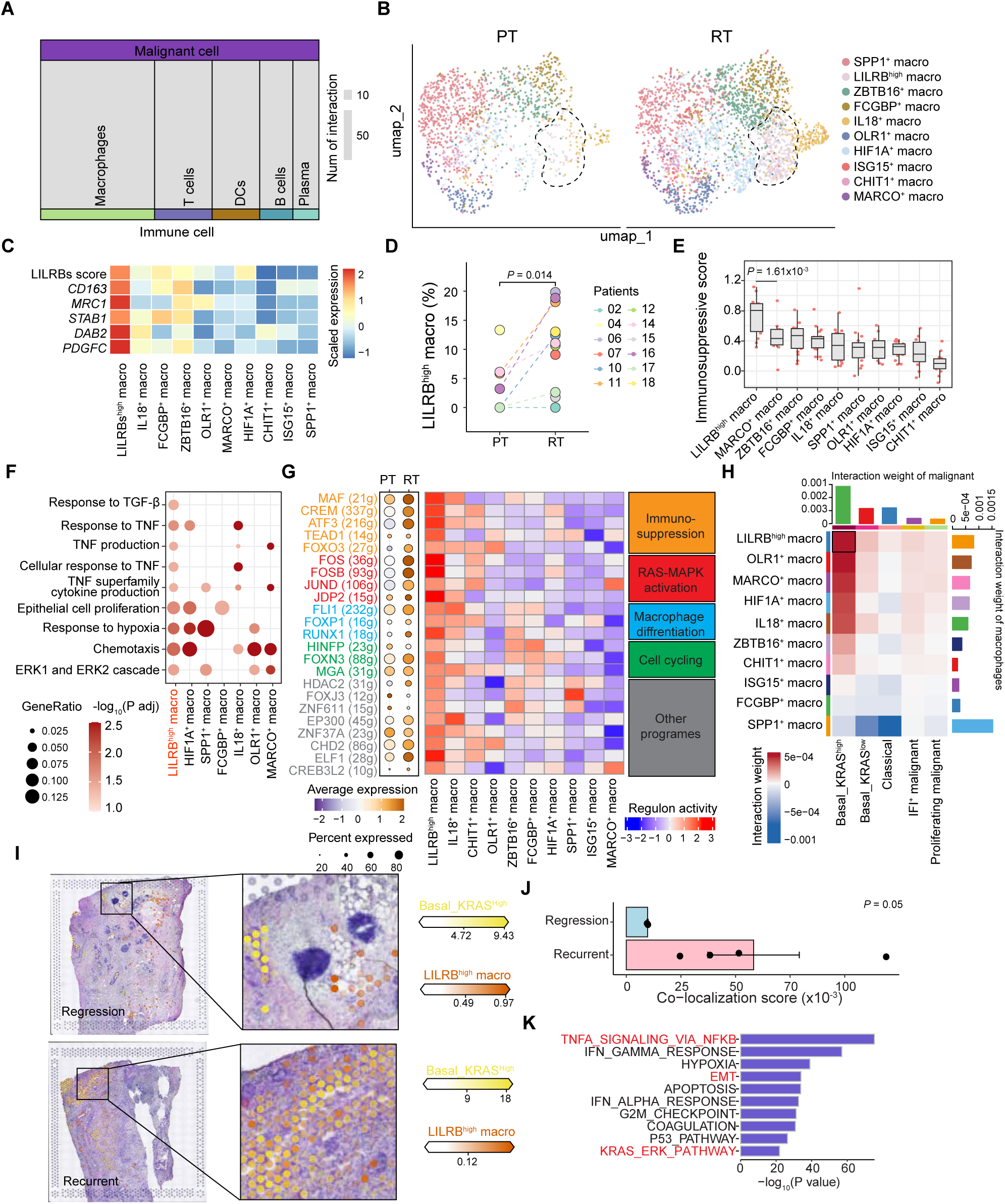
Immunosuppressive LILRB^high^ macrophages expand in recurrent PDAC and preferentially interact with Basal_KRAS^high^ malignant cells. **(A)** LIANA-ranked ligand–receptor interaction landscape across major cell compartments, showing the number of predicted interactions between malignant cells and immune lineages. **(B)** UMAP of snRNA-seq macrophages from matched primary tumors (PT) and recurrent tumors (RT), annotated into macrophage subtypes; LILRB^high^ macrophages are highlighted. **(C)** Heatmap of LILRB score and representative TAM markers across macrophage subtypes. **(D)** Paired quantification of LILRB^high^ macrophage fraction among total macrophages in PT versus RT across patients; lines connect matched samples; *P* value from t-test. **(E)** Immunosuppressive score across macrophage subtypes; boxes show median and interquartile range with whiskers indicating 1.5× IQR; points denote individual samples/cells; *P* value from group comparison between LILRB^high^ macro and MARCO^+^ macro. **(F)** Pathway enrichment across macrophage subtypes. Dot size indicates gene ratio and color denotes significance. **(G)** SCENIC regulon analysis of macrophage subtypes showing transcription factor regulon expression level dot size represents the fraction of regulon-positive cells, dot color represents the expression level. Heatmap color represents the regulon activity. The color of regulon text represents their biological function. **(H)** CellChat-derived interaction weights between malignant states (columns) and macrophage subtypes (rows). Heatmap color represents the interaction strength level. Side bars summarize total interaction weight for each malignant state and macrophage subtype. **(I)** Spatial mapping of Basal_KRAS^high^ malignant cells and LILRB^high^ macrophages in representative regression and recurrent samples, with zoomed-in regions indicated. **(J)** Quantification of Basal_KRAS^high^–LILRB^high^ spatial co-localization score in regression versus recurrent samples. *P* value, one-sided t-test. **(K)** Enriched Hallmark pathways associated with the Basal_KRAS^high^–LILRB^high^ module; bars show −log_10_ (Fisher *P* value). PT, primary tumor; RT, recurrent tumor; TAM, tumor-associated macrophage; LILRB^high^, macrophages with high *LILRB1–LILRB5* expression; SCENIC, single-cell regulatory network inference and clustering. See also **Figs. S4–S6**.

Functionally, these macrophages were enriched for TGF-β, TNF-α and epithelial proliferation pathways, as well as small GTPase-related signaling (**Fig. 4F** and **Fig. S4E**). Regulatory network analysis using SCENIC^30^ revealed elevated activity of immunoregulatory transcription factors (*MAF*, *CREM,* and *ATF3*) in LILRB^high^ macrophages, with further increases in recurrence (**Fig. 4G** and **Fig. S4F**). Regulons linked to RAS signaling (*FOS*, *JUND*), macrophage differentiation (*RUNX1*) and proliferation were also activated.

Notably, Basal_KRAS^high^ tumor cells exhibited the strongest interactions by CellChat^31^ with LILRB^high^ macrophages (**Fig. 4H**). Correlation analyses revealed a tightly coupled small module comprising Basal_KRAS^high^ cells, Tregs, and LILRB^high^ macrophages (**Fig. S5A**). Spatial transcriptomics further confirmed preferential co-localization of Basal_KRAS^high^ tumor cells and LILRB^high^ macrophages, which was significantly enhanced in recurrent tumors (**Fig. 4I, J** and **Fig. S5B**).

To elucidate the functional consequences of Basal_KRAS^high^–LILRBs^high^ macrophage crosstalk, we interrogated signaling pathways associated with their interactions. Pathway analysis revealed increased activity of MHC-II, TGF-β, MK, and FN1 signaling in recurrent tumors (**Fig. S5C, D**), pathways previously implicated in tumor progression^32–36^. Using NicheNet^37^ to prioritize ligand–receptor interactions, we identified *HLA-A/B/C, MDK, TGFBI* and *FN1* as top-ranked ligands expressed by Basal_KRAS^high^ cells in recurrence, with corresponding receptors (*LILRB1/2*, *NOTCH2,* and integrins) on LILRB^high^ macrophages (**Fig. S5E–G**). These predicted interactions converged on activation of TNFα, EMT and KRAS–ERK signaling programs (**Fig. 4K**) and were linked to transcriptional regulators such as *ATF3, FOS, JUND* and *MAF*, supporting expansion and immunosuppressive polarization of LILRB^high^ macrophages (**Fig. S5F**).

Collectively, these findings delineate a KRAS-activated tumor–macrophage circuit in which Basal_KRAS^high^ cells spatially and functionally engage LILRB^high^ macrophages, establishing an immunosuppressive niche that characterizes recurrent PDAC.

### LILRB4^+^ macrophages drive recurrent PDAC

We next sought to identify which LILRB family members contribute most to recurrence. Among the five LILRB genes, only *LILRB4* was significantly upregulated in macrophages from recurrent tumors (**Fig. 5A**). Within the LILRB^high^ subset, *LILRB4* expression alone was specifically enriched in recurrence (**Fig. 5B**). Given prior reports linking LILRB4^+^ macrophages to pro-tumorigenic functions across cancer types^38–41^, we prioritized *LILRB4* for further investigation as a candidate biomarker and therapeutic target.

**Figure 5.**
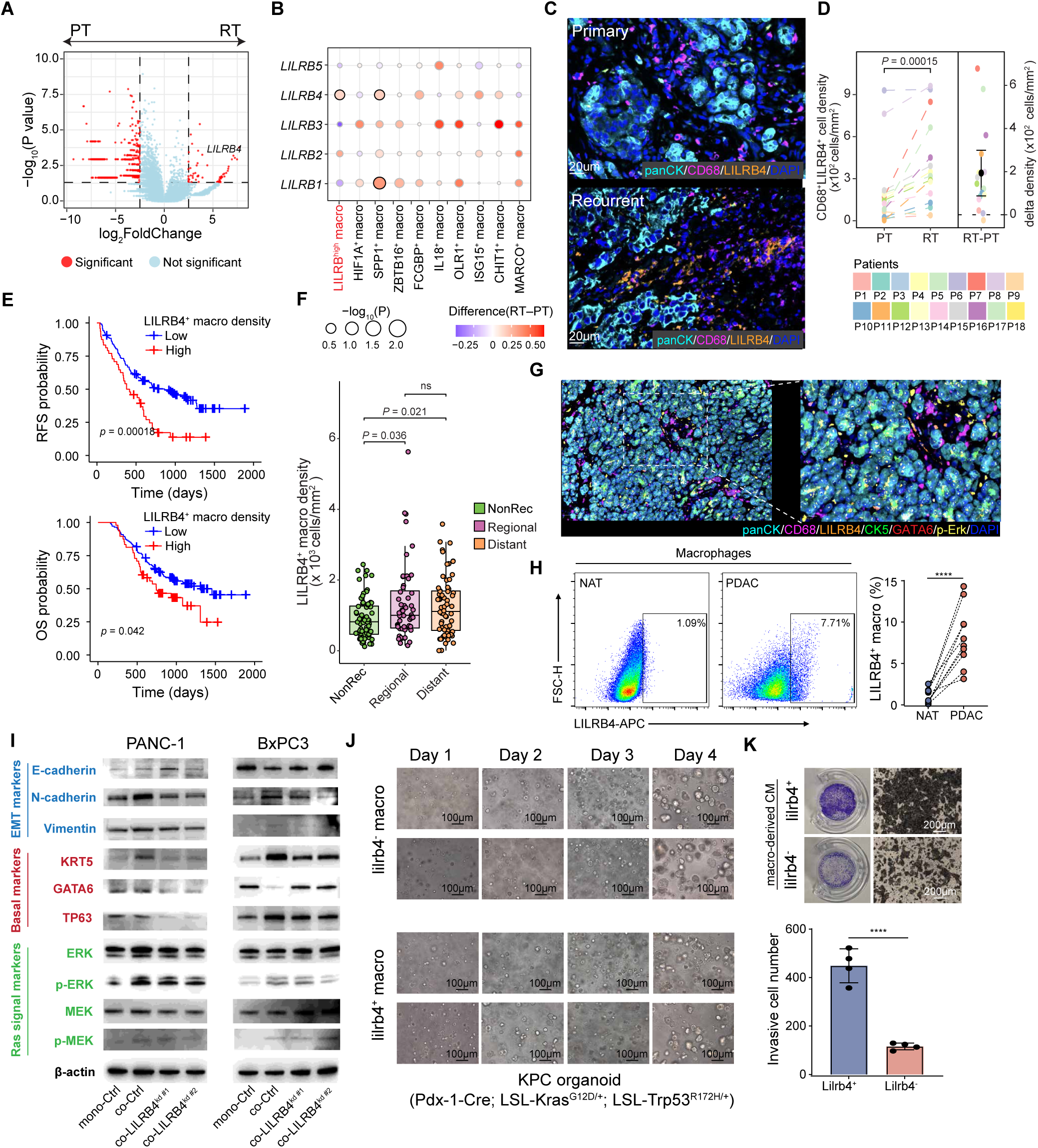
LILRB4 marks recurrence-associated LILRB^high^ macrophages and couples with KRAS-activated basal malignant cells to promote aggressive disease. **(A)** Volcano plot of differential expressed genes in macrophages between primary tumors (PT) and matched recurrent tumors (RT), highlighting *LILRB4* as a recurrence-associated marker. **(B)** Dot plot of LILRB family genes’ expression across macrophage subtypes. Dot size indicates *P* value significance and color denotes expression change between RT and PT (RT–PT), identifying *LILRB4* as selectively increased within LILRB⁺ macrophages. **(C)** Representative multiplex immunofluorescence (mIF) images showing LILRB4^+^CD68^+^ macrophage distribution in matched PT and RT tissues; stains include LILRB4 (orange), CD68 (magenta), PanCK (light blue), and DAPI (blue). Scale bars, 100 µm. **(D)** Quantification of LILRB4^+^CD68^+^ macrophage density in matched PT and RT samples (left) and paired change (RT–PT) (right); lines connect matched samples; *P* value from paired t-test. **(E)** Kaplan–Meier analyses of progression-free/disease-free survival (PFS/DFS; top) and overall survival (OS; bottom) stratified by intratumoral LILRB4^+^ macrophage density (High vs Low, separated by top quarter) in an independent 190 patient’s cohort. *P* values, log-rank test. **(F)** LILRB4^+^ macrophage density across clinical recurrence categories (non-recurrent, regional recurrence, and distant metastasis); boxes show median and interquartile range with whiskers indicating 1.5× IQR; significance is indicated (*P* value, Wilcoxon test; ns, not significant). **(G)** Representative mIF images showing spatial co-localization of LILRB4^+^ macrophages (LILRB4^+^ CD68^+^) with Basal_KRAS^high^ (panCK^+^ CK5^+^ p-Erk^+^ GATA6^-^) tumor cells in matched PT and RT tissues; scale bars, 100 µm. **(H)** Representative flow cytometry plots (left) showing LILRB4 expression in tumor-infiltrating macrophages from matched non-tumor adjacent tissue (NAT) and PDAC tumor tissue. LILRB4⁺ macrophages were rare in NAT and enriched in PDAC. Right, paired quantification of LILRB4^+^ macrophage frequency (%) of all macrophages in NAT versus PDAC across samples (****, *P* < 10^-4^; paired test). **(I)** Immunoblot analysis of PANC-1 and BxPC-3 cells under monoculture or co-culture with TAMs, including control TAMs and LILRB4-knockdown TAMs (LILRB4^kd#1^ and LILRB4^kd#2^). Blots show epithelial/plasticity and lineage markers (E-cadherin, N-cadherin, Vimentin, KRT5, GATA6, TP63) and ERK–MEK pathway activation (ERK, p-ERK, MEK, p-MEK); β-actin, loading control. **(J)** Representative bright-field time-course images (Days 1–4) of KPC PDAC organoids (Pdx1-Cre; LSL-Kras^G12D/+;^ LSL-Trp53^R172H/+^) co-cultured with LILRB4⁻ versus LILRB4⁺ TAMs, showing enhanced organoid growth, morphologic plasticity, and invasive outgrowth in the LILRB4⁺ TAM condition. **(K)** Representative transwell invasion assay images and quantification of invaded PDAC cells after treatment with TAM-derived conditioned medium (CM) from LILRB4⁺ or LILRB4⁻ TAMs. LILRB4⁺ TAM-derived CM increased tumor-cell invasion (*** *P* < 10^-4^). PT, primary tumor; RT, recurrent tumor; TAM, tumor-associated macrophage; mIF, multiplex immunofluorescence; PanCK, pan-cytokeratin; PFS/DFS, progression-free/disease-free survival; OS, overall survival. TAM, tumor-associated macrophage; NAT, non-tumor adjacent tissue; CM, conditioned medium. See also **Fig. S6**.

The mIF analysis of an independent validation cohort confirmed a marked increase in LILRB4^+^ macrophages in recurrent versus matched primary tumors (**Fig. 5C, D**). In the 190-patient whole-slide mIF cohort, high LILRB4^+^ macrophage density was associated with worse relapse-free survival (*P* = 1.8 × 10⁻⁴) and overall survival (*P* = 0.042) (**Fig. 5E** and **Fig. S6A**). *LILRB4*^+^ macrophages were also enriched in patients with local recurrence and distant metastasis (**Fig. 5F**). Spatially, LILRB4^+^ macrophages co-localized with p-ERK^+^CK5^+^GATA6^−^ tumor cells, consistent with direct interaction with KRAS-activated basal-like cells (**Fig. 5G**).

Flow cytometry demonstrated that LILRB4^+^ macrophages were rare in adjacent non-tumor tissue but enriched within PDAC tumors (**Fig. 5H** and **Fig. S6B**), supporting a tumor-specific macrophage program. In co-culture assays, LILRB4^+^ macrophages induced basal-like and plasticity-associated transcriptional changes in two human PDAC cell lines (Panc-1, harboring a *KRAS*_G12D mutation, a model of basal-like PDAC, the *KRAS* wild-type BxPC-3 cell line, a representative of the classical PDAC) and enhanced ERK and MEK phosphorylation. Knockdown of LILRB4 in macrophages attenuated these effects (**Fig. 5I, Fig. S6C-E**). Similar findings were observed in murine systems: LILRB4^+^ macrophages isolated from orthotopic KPC tumors promoted basal-like morphological reprogramming and increased invasiveness of KPC-derived organoids compared with LILRB4^−^ macrophages (**Fig. 5J, K**).

Collectively, these results identify LILRB4 as a recurrence-associated macrophage effector that enhances KRAS–ERK signaling, tumor cell plasticity and invasive behavior, establishing LILRB4^+^ macrophages as a functional driver of the immunosuppressive niche in recurrent PDAC.

### Co-targeting KRAS and LILRB4 produces superior tumor control

To evaluate the therapeutic potential of targeting LILRB4, we established a PDO–macrophage co-culture system. Importantly, we developed hu36B10D1, a first-in-class human LILRB4-targeting monoclonal antibody (mAb). Treatment with anti-LILRB4 mAb significantly reduced organoid growth and tumor cell viability compared with isotype controls (**Fig. 6A**), indicating that LILRB4 blockade suppresses macrophage-driven tumor support. Conditioned medium from LILRB4 mAb treated co-cultures similarly reduced tumor cell invasion in transwell assays, consistent with on-target inhibition of macrophage-derived pro-invasive signals (**Fig. 6B**). Notably, using a Luc-PANC1 reporter system, LILRB4 blockade had minimal effect on tumors formed by PANC-1 cells alone but markedly suppressed tumor growth in macrophage-containing xenografts (**Fig. 6C**), confirming macrophage-dependent therapeutic activity.

**Figure 6.**
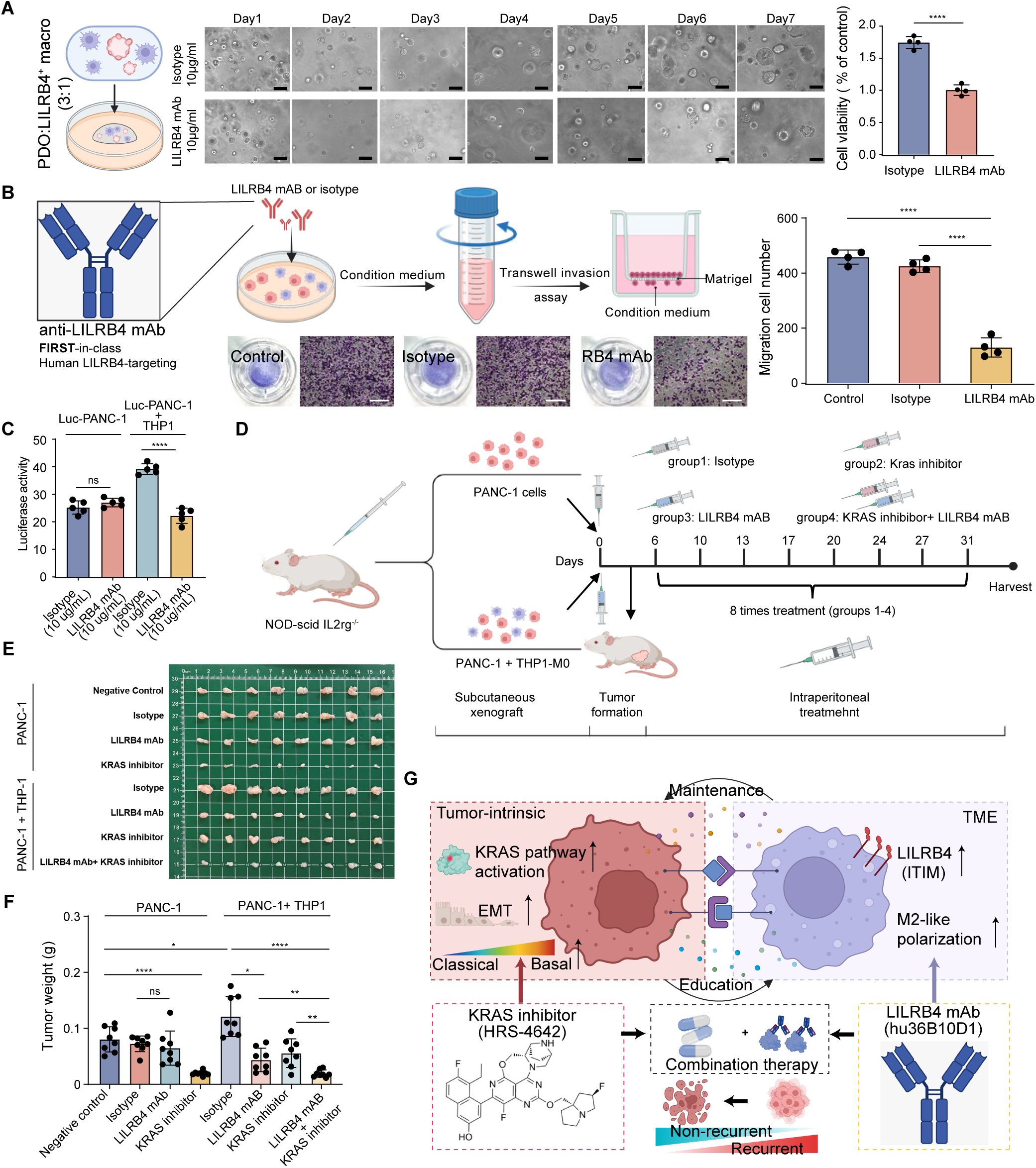
Dual KRAS and LILRB4 targeting produces superior tumor control in macrophage-supported PDAC. **(A)** Patient-derived organoid (PDO)–THP-1 co-culture assay (PDO: THP-1 = 3:1) treated with isotype control or LILRB4 monoclonal antibody (mAb; 10 μg mL⁻¹). Representative bright-field images are shown over 7 days (left), and CellTiter-Glo quantification of tumor-cell viability is shown at right. (**** *P* < 10^-4^). **(B)** Schematic of hu36B10D1 (human anti-LILRB4 mAb, Nanjing Leads Biolabs Co., Ltd.) and conditioned-medium workflow for transwell invasion assays. Conditioned medium was collected from tumor–macrophage co-cultures treated with control, isotype, or anti-LILRB4 mAB and applied to Matrigel-coated transwells. Representative invasion images and quantification (right) of migrated/invaded cells are shown. (**** *P* < 10^-4^). **(C)** Luciferase activity in xenografts generated from Luc-PANC-1 cells alone or Luc-PANC-1 + THP-1 co-inoculation, with isotype or LILRB4 mAb treatment (10 μg mL⁻¹), showing macrophage-dependent antitumor effects of LILRB4 blockade. (**** *P* < 10^-4^, ns, not significant). **(D)** Experimental design for *in vivo* LILRB4 blockade in a macrophage-containing xenograft model. THP-1 and PANC-1 cells were co-inoculated subcutaneously into NOD-scid Il2rg^-/-^ mice, followed by intraperitoneal treatment with isotype or anti-LILRB4 mAb at the indicated schedule. **(E)** Representative excised tumors from PANC-1–only and PANC-1 + THP-1 xenograft models treated with negative control, isotype, LILRB4 mAb, KRAS inhibitor, or the combination (LILRB4 mAb + KRAS inhibitor), with enlarged views of representative tumors. **(F)** Tumor weight quantification for treatment groups shown in (**E**). In the macrophage-containing model (PANC-1 + THP-1), combined LILRB4 mAb and KRAS inhibitor treatment produced the strongest tumor suppression. (**** *P* < 10^-4^, ** *P* < 0.01, * *P* < 0.05, ns, not significant). **(G)** Working model summarizing reciprocal tumor–immune reinforcement in recurrent PDAC: KRAS activation and EMT-associated plasticity promote a basal-like malignant state, while LILRB4^+^ tumor-infiltrating macrophages maintain an immunosuppressive niche. Combined KRAS inhibition (HRS-4642) and LILRB4 blockade (hu36B10D1) disrupt this bidirectional circuit and suppress recurrence-associated tumor programs. PDO, patient-derived organoid; mAb, monoclonal antibody; TAM, tumor-associated macrophage; TME, tumor microenvironment. See also **Fig. S7**.

Given the expansion of Basal_KRAS^high^ cells in recurrence, we next examined whether KRAS inhibition modulates this tumor–immune axis. We established an immunocompetent KPC GEMM syngeneic allograft with acquired resistance to KRAS inhibition by transplanting KRAS inhibitor–resistant KPC tumor cells into syngeneic recipient mice. Following KRAS inhibitor treatment (HRS-4642), the murine tumor cell cluster corresponding to human Basal_KRAS^high^ cells showed a marked reduction in abundance (**Fig. S7A–C**). In contrast, infiltration of the macrophage cluster corresponding to human *LILRB4*⁺ macrophages remained unchanged between treatment and control groups (**Fig. S7D–F**). These findings indicate that KRAS inhibition selectively suppresses KRAS-activated basal tumor cells but fails to eliminate LILRB4⁺ macrophage infiltration, thereby supporting combined KRAS inhibitor and LILRB4 monoclonal antibody therapy as a strategy to overcome macrophage-associated resistance.

We then evaluated whether LILRB4 blockade enhances the efficacy of KRAS inhibition in PDAC. In subcutaneous implantation of PANC-1 cells without macrophages, KRAS inhibitor treatment significantly suppressed tumor growth, whereas LILRB4 mAb alone had minimal antitumor effect. In contrast, in macrophage-containing xenografts generated by co-injecting PANC-1 cells with macrophages, both KRAS inhibitor and LILRB4 mAb reduced tumor growth; however, the effect of KRAS inhibition was attenuated compared with the tumor-only setting, consistent with macrophage-mediated resistance. Strikingly, combined treatment with KRAS inhibitor and LILRB4 mAb achieved the most pronounced tumor suppression in the macrophage-containing model, outperforming either monotherapy (**Fig. 6E, F**).

Together, these data support a model in which KRAS activation and EMT-related plasticity in tumor cells promote a classical-to-basal shift, while LILRB4^+^ macrophages maintain an M2-like immunosuppressive state in the TME. These tumor-intrinsic and microenvironmental programs reinforce each other through bidirectional signaling, and dual treatment with KRAS inhibitor (HRS-4642) plus LILRB4 mAb interrupts this loop and shifts tumors toward a less recurrence-prone state (**Fig. 6G**).

## Discussion

Despite advances in surveillance and systemic therapy, the prognosis of PDAC remains poor, largely owing to the high incidence of postoperative recurrence. Recurrent tumors are typically managed based on the molecular features of the primary lesion, yet whether they preserve—or remodel—their TME during relapse has remained unclear. A recent study demonstrated that progressive selective pressures during invasion and metastasis reshape PDAC phenotypes, showing that variability in KRAS genotype dependence and BRCA2 inactivation between primary and recurrent tumors influences therapeutic response and evolutionary trajectories^42^. However, these analyses have largely emphasized tumor-intrinsic alterations. By generating a longitudinal single-cell atlas of matched primary and recurrent PDAC, we delineate how oncogenic signaling and immune remodeling converge to shape relapse.

Consistent with prior genomic profiling, *KRAS* mutant allele dosage was maintained or amplified in recurrent tumors, reinforcing its trunk-driver role in PDAC evolution. Yet our data indicate that genetic persistence alone does not fully account for the aggressive phenotype of recurrence. Instead, we observe enrichment of a basal-like, pEMT program specifically in recurrent lesions. Unlike cross-sectional comparisons between primary and metastatic sites, our paired, same-organ design minimizes anatomical confounders and attributes this basal-pEMT shift to temporal evolution under postoperative and immune-mediated selection pressures. Sustained Ras activation in this context appears to promote lineage plasticity rather than simply proliferation, enabling residual tumor cells to adapt and repopulate the remnant pancreas.

A central conceptual advance of this study is the identification of LILRB4^+^ TAMs as active participants in this plastic transition. Although myeloid-mediated immunosuppression in PDAC is well established, the mechanisms by which macrophages influence tumor cell differentiation have remained poorly defined. Our findings position LILRB4^+^ TAMs as drivers of basal–EMT reprogramming, bridging tumor-intrinsic KRAS activation with extrinsic immune cues. Spatial and functional analyses support a bidirectional circuit in which KRAS-activated basal tumor cells engage LILRB4^+^ macrophages to create a niche that protects residual cells and reinforces aggressive lineage states. This model reframes recurrence as a cooperative process between tumor cells and the immune microenvironment rather than a purely cell-autonomous evolutionary event. Translationally, our data suggest that Ras-directed therapies alone may be insufficient when a pro-plasticity myeloid niche is established. While KRAS-targeted vaccines and inhibitors show early promise, their durability is limited by heterogeneous Ras dependence and profound immunosuppression. We demonstrate that LILRB4^+^ TAM infiltration is associated with adverse clinical outcomes and that LILRB4 blockade enhances the efficacy of KRAS inhibition in macrophage-containing models. These findings support a therapeutic paradigm in which dismantling the tumor–myeloid axis enhances susceptibility to oncogenic pathway inhibition. Co-targeting KRAS and LILRB4 may therefore both relieve immune suppression and disrupt macrophage-driven lineage plasticity, providing a rational combinatorial strategy to prevent or delay recurrence.

This study is strengthened by a rare longitudinal cohort of matched primary and intrapancreatic recurrent tumors, enabling direct interrogation of temporal tumor–immune evolution within the same anatomical context. Integration of multi-omics profiling with spatial validation and animal models further links molecular insights to clinical relevance. Our macrophage-containing mouse model has inherent limitations and does not fully capture the complexity of the human PDAC microenvironment. Validation in more biologically relevant systems will be necessary to confirm the generalizability of our findings. Nevertheless, LILRB4 blockade shows promise in combination with KRAS inhibition for recurrent tumors. The efficacy, durability, and safety of this combinatorial strategy warrant further rigorous evaluation prior to clinical translation. Future work should also delineate the specific tumor-derived ligands and metabolic cues that activate LILRB4 signaling, as well as the transcriptional circuitry coupling Ras signaling to basal-pEMT reprogramming under myeloid influence.

In summary, our findings redefine PDAC recurrence as a dynamic ecosystemal process driven by convergence of intrinsic KRAS-mediated plasticity and extrinsic macrophage remodeling. Targeting this KRAS–LILRB4 tumor–immune circuit offers a framework for next-generation combination strategies aimed at improving the durability of response in this lethal disease.

## Online Methods

### Ethics statement

Written informed consent was obtained from all patients whose clinical information and tissues were used. The study was conducted in accordance with recognized ethical guidelines of Declaration of Helsinki and approved by the Institutional Review Boards of the First Affiliated Hospital of Nanjing Medical University (Nanjing, Jiangsu Province, China; 2025-SR-338).

### Study cohorts

For the cohort for survival analysis, clinical information of 2710 patients were collected between December 2014 and December 2024. Samples from patients with pancreatic cancer were prospectively collected via the Pancreas Biobank of the First Affiliated Hospital of Nanjing Medical University, which is a part of Jiangsu Biobank of Clinical Resource. Following discharge, patients underwent regular follow-up visits every 3 months, with tumor recurrence monitoring performed every 2 to 3 months. For the discovery cohort, longitudinal frozen-free samples with paired tumor, adjacent normal tissues and peripheral blood were collected from 18 treatment-naive primary or post-chemotherapy recurrent PDAC patients. Table S2 shows detailed clinical and pathological information including age, gender, tumor stage, nodal status, prognosis. The external validation cohort including formalin-fixed paraffin-embedded (FFPE) tissue blocks from 190 pathologically diagnosed with PDAC patients who underwent radical-intent surgery for pancreatic cancer at Pancreas Center of the First Affiliated Hospital of Nanjing Medical University.

### Sample processing

The fresh-frozen tumor tissues were snap-frozen in liquid nitrogen immediately after surgical resection and subsequently stored at −80°C. FFPE tumor tissues were obtained from archived residual clinical specimens. Blood samples were collected prior to treatment and stored at −80°C.

### Single nuclei isolation and sequencing

To isolate single nuclei from frozen tissue, begin by cutting the tissue into small pieces (1-2 mm³) and place it in 3 mL of lysis buffer (10 mM Tris, pH 7.4; 10 mM NaCl; 3 mM MgCl₂; 0.05% NP-40 detergent). Homogenize the tissue using a glass Dounce homogenizer, first with a loose pestle for 10 strokes, followed by 5 strokes with the tight pestle. Allow the sample to lyse for 5 minutes on ice, then add 5 mL of wash buffer (10 mM Tris, pH 7.4; 10 mM NaCl; 3 mM MgCl₂; 1% BSA; 1 mM DTT; 1 U/µL RNase inhibitor; nuclease-free water). Pass the lysate through a 30-μm cell strainer to remove large debris and centrifuge at 500×g for 5 minutes. After pelleting, resuspend the nuclei in 5-10 mL of wash buffer by pipetting up and down 8-10 times. Wash the nuclei 3 times by centrifuging and resuspending them in fresh wash buffer. After the final wash, resuspend the nuclei in 1 mL of wash buffer, mix with 25% Optiprep, and layer the sample on top of a 29% Optiprep cushion. Perform density gradient centrifugation at 10,000×g for 30 minutes. After centrifugation, carefully collect the nuclear layer, resuspend the nuclei in wash buffer, and wash 3 more times. To assess nuclear quality, count the nuclei and observe their morphology under a microscope using AO/PI staining. Finally, resuspend the nuclei in Nuclei Resuspension Buffer (1X 10X Genomics Nuclei Buffer, 1 mM DTT, 1 U/µL RNase inhibitor, nuclease-free water) to a final concentration of approximately 1×10⁶ nuclei/mL. Finally, the nucleus suspension were processed with the 10x Chromium Single Cell 3’ Kit as per the manufacturer’s instructions at Novogene Bioinformatics Technology Co., Ltd (Tianjin, China).

The nucleus suspension was loaded into 10x Chromium Chip and barcoded with a 10x Chromium Controller. RNA from the barcoded nucleus was subsequently reverse-transcribed, amplified and prepared into sequencing libraries with 10x Library Construction Kit according to the manufacturer’s instructions. Sequencing was performed with Illumina NovaSeq with 150-bp paired-end reads at Novogene Bioinformatics Technology Co., Ltd (Tianjin, China).

### WES data analysis

Read pairs in FASTQ format were trimmed and filtered for quality using fastq-mcf (https://github.com/ExpressionAnalysis/ea-utils). High-quality reads were aligned to the human reference genome (GRCh38) using the Burrows-Wheeler Aligner (BWA 0.7.12). The resulting BAM files were processed using the Genome Analysis Toolkit (GATK) to enhance alignment accuracy. Key steps included marking duplicates, performing local realignment around high-confidence insertions and deletions (INDELs), and recalibrating base quality scores. Somatic point mutations were identified using Strelk2^43^. A multi-step filtering process was applied to ensure high-quality mutation calls, including criteria such as minimum read depth, strand-bias checks, and exclusion of artifacts using a panel of normals^44^. This pipeline ensured a robust and reliable set of somatic mutation data for downstream analyses.

For the calculation of tumor VAF, we converted the Strelka output VCF to a tabular format and extracted allele counts. We then used *ICAMS*^45^ (function “MakeDataFrameFromVCF” and “GetStrelkaVAF”) to calculate VAF for each variant based on the reference and alternate read counts. For established PDAC driver genes, we generated per-sample MAF files (*vcf2maf*^46^) and loaded them with *maftools*^47^. We selected variants by Hugo_Symbol for genes of interest and matched these selected variants back to the VAF tables by genomic position and retained the VAF values for downstream analyses.

We estimated mutation allele dosage by integrating VAF, allele-specific copy number, and tumor purity. We first computed VAF for each somatic variant, then annotated each mutation with the local total copy number by intersecting the mutation genomic coordinate with absolute copy-number segments. For each mutation, we assigned the segment “Modal_Total_CN” as the total copy number (*CN*_total_) at that locus. We also obtained tumor purity (*ρ*) and matched purity values to samples. Finally, we calculated mutation dosage as a purity-adjusted copy-number–scaled measure of mutant allele signal:

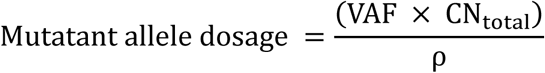

We excluded variants with missing copy-number or purity annotations before downstream analyses.

### Bulk RNAseq data analysis

For each sample, RNA-seq clean reads were obtained and aligned to the GRCh38 reference genome (ftp://ftp.ensemblgenomes.org/) using *HISAT2*^48^. Sequencing read counts were calculated using *Stringtie*^49^ and expression levels were normalized across samples using the Trimmed Mean of M values (TMM) method. Normalized expression levels were converted to FPKM (Fragments Per Kilobase of transcript per Million mapped fragments). Differential gene expression analysis was performed using the R package *edgeR*^50^. P-values were calculated, and multiple hypothesis testing was controlled by the Benjamini-Hochberg algorithm to calculate the False Discovery Rate (FDR), with corrected p-values referred to as q-values.

Basal-like and classical PDAC subtype scores were computed from normalized bulk RNA-seq expression matrices using Gene Set Variation Analysis (GSVA). For each sample, GSVA enrichment scores were calculated for a basal-like gene set (*S*_basal_) and a classical gene set (*S*_classical_)^10^. We defined a subtype axis score as:

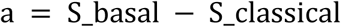

To obtain a bounded 0–1 metric for visualization, we converted the axis score to “basallike” using a logistic transformation:

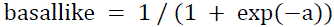

Basallike approaches 1 for basal-like samples and 0 for classical-like samples. For paired PT) and RT samples, the within-patient shift was summarized as:

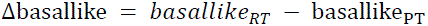

Gene set function enrichment analysis was performed using the R package *fgsea*, with enrichment criteria set to q-value < 0.05. Heatmaps of specific genes were generated using the R package *pheatmap.* Gene-set signature scores were computed for each sample by averaging normalized expression values across the genes in each predefined gene sets.

### snRNAseq data preprocessing and clustering

Raw reads from snRNA-seq were processed using the Cell Ranger pipeline (version 6.0.0, 10× Genomics) and mapped to the GRCh38 reference genome to generate gene count matrices indexed by cell barcodes. The resulting gene-barcode matrices were analyzed using the *Seurat*^51^ R package. Cells expressing at least 200 genes and genes detected in at least 3 cells were retained for further analysis. All samples were merged into a single Seurat object and filtered based on certian quality control criteria. These preprocessing steps ensured the retention of high-quality cells for downstream analyses.

Clustering and dimension reduction were performed on the filtered Seurat object, which was normalized and scaled with the mitochondrial gene percentage regressed out. The top 2000 most variable genes were identified using the “FindVariableFeatures” function and used for principal component analysis (PCA). Nearest neighbors were identified using the “FindNeighbors” function, and graph-based clustering was performed with “FindCluster” to define cell subtypes. The uniform manifold approximation and projection (UMAP) algorithm was applied for cell subtype visualization. To correct for batch effects, the harmony algorithm (R package *Harmony*^14^) was applied before clustering. Cells were first partitioned into broad categories then further subclustered into subtypes for each category. Cluster identities were assigned based on marker gene expression and manually reviewed to ensure accurate cell type annotation. DEGs among clusters were identified using the “FindAllMarkers” function. The KRAS pathway score was calculated using KRAS pathway genes from the gene sets downloaded from the Molecular Signatures Database^52^ (MsigDB, https://www.gsea-msigdb.org/gsea/msigdb/). The functional enrichment analysis were conducted by R package *clusterProfier*^53^.

### Spatial transcriptomics datasets deconvolution and co-localization analysis

To improve the mapping of scRNA-seq clusters to spatial spots, the cellular composition of each spatial spot was deconvoluted using the cell2location Python module, following the tutorial guidelines from the Cell2location website. The single-cell regression model was trained with parameters max_epochs=250 and lr=0.002, while the final cell2location model was trained with parameters max_epochs=30,000. This integration provided a detailed spatial map of cellular compositions.

We quantified spatial co-localization between Basal_KRAS^high^ tumor signal and LILRBs⁺ macrophage signal using spot-level metadata from each Visium Seurat object. We converted both features from percentages to proportions (0–1) by dividing by 100.

For each sample, we calculated two complementary co-localization metrics across tissue spots. First, we computed a product-based co-localization score as the mean of the per-spot product of the two proportions:

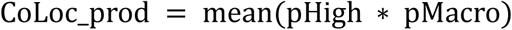

We then compared per-sample co-localization scores between relapse and regression groups.

### Trajectory and cell differentiation analysis

Trajectory analysis was performed using *scVelo*^54^ by integrating spliced and unspliced transcript counts with the annotated single-cell expression matrix. After merging, genes were filtered and normalized, and then first/second-order moments were computed with “scv.pp.moments”. RNA velocity vectors were then estimated with “scv.tl.velocity”, and the cell–cell transition graph was constructed using “scv.tl.velocity_graph”, with velocities projected onto the UMAP embedding for visualization. To infer a continuous trajectory, we fit the dynamical model using scv.tl.recover and computed latent time with “scv.tl.latent_time”, which was used as pseudotime to order cells along the inferred trajectory.

*CytoTRACE2* ^17^was applied to malignant tumor cells from RT samples to infer differentiation potential. After removing predicted doublets, *CytoTRACE2* was run on the Seurat object using raw RNA counts as input and resulting *CytoTRACE2* scores were added to the cell metadata and visualized on the UMAP embedding. For sample-level comparisons, *CytoTRACE2* scores were averaged within each tumor state per sample. An ordered trend across states was assessed using a linear mixed-effects model with sample as a random effect.

### Scenic analysis

*SCENIC*^30^ was performed on the macrophage single-cell dataset to infer transcription factor (TF) regulons and their cell-state–specific activities. Regulon activities were summarized at the cell-type level by averaging AUC scores across cells, and cell-type specificity was assessed using the Regulon Specificity Score (RSS); AUCell-based binarization was additionally used to estimate the fraction of cells with active regulons. Finally, regulon AUC matrices were integrated back into the Seurat metadata for visualization and downstream comparisons across macrophage subtypes and conditions.

### Cell-cell communication analysis

Cell–cell communication analysis was performed using *LIANA*^21^ on a normalized Seurat object containing RT malignant and immune cell populations. Ligand–receptor interactions were inferred and summarized by consensus ranking, and high-confidence interactions were retained using an aggregate-rank cutoff (aggregate_rank ≤ 0.3) with unique source–target–ligand–receptor pairs. The number of tumor–immune interactions was then quantified in both directions and visualized as an alluvial diagram of interaction counts by partner cell type.

Next, we used *CellChat* ^31^to analyze cell-to-cell communication between macrophages and tumor cells. A CellChat object was created by grouping the cells into predefined clusters. The ligand-receptor interaction database used for this analysis was “CellChatDB.human”, without additional supplementation. Preprocessing steps were conducted using default parameters provided by the package. To infer the communication network, the functions “computeCommunProb” and “computeCommunProbPathway” were applied for each ligand-receptor pair and each signaling pathway, separately.

Finally, we used *NicheNet*^37^ to infer the interaction mechanisms between Basal_KRAS^high^ cells (sender cells) and LILRBs^high^ macrophages (affected cells) in the scRNAseq dataset. For ligand-receptor interactions, only clustered cells with gene expression levels above 10% were considered. The RT group served as the condition of interest for prioritizing ligands. The top 300 ligands and top 2000 target genes from differentially expressed genes of the sender and affected cells were extracted for paired ligand-receptor activity analysis. To assess functional gene sets enriched by top-ranked ligands, gene sets downloaded from MsigDB^99^, were used as potential targets. Functional enrichment was cross-validated using the Fisher’s exact test in two rounds, and the average p-value was calculated to ensure robust prioritization of ligand-receptor interactions and their functional implications.

### Multiplex immunofluorescent staining

FFPE tissue sections (5μm thick) were deparaffinized in xylene and rehydrated through a graded ethanol series. Multiplex immunofluorescence staining was performed using the 7-color Panovue Multiplex Immunohistochemistry Kit (Cat# 10268100100) according to the manufacturer’s instructions. Briefly, after rehydration, slides were subjected to heat-induced epitope retrieval in EDTA buffer (pH 9.0) using a microwave. Following a 10-minute blocking step, the slides were incubated with the primary antibody for 1 hour and then with the secondary antibody for 30 minutes. After each antibody incubation, TSA (tyramide signal amplification) labeling was applied, followed by antibody stripping through antigen retrieval before the next staining cycle. Finally, nuclei were counterstained with DAPI.

### Cell culture

The human pancreatic cancer cell lines (BxPC-3 and PANC-1) and the human monocytic cell line THP-1 were obtained from Shanghai Cell Bank (Shanghai, China). BxPC-3 and THP-1 cells were cultured in RPMI-1640 medium, whereas PANC-1 cells were maintained in Dulbecco’s Modified Eagle Medium (DMEM). All culture media were supplemented with 10% fetal bovine serum (Wisent, Canada) and 1% penicillin-streptomycin (MCE, USA). To induce macrophage differentiation, THP-1 monocytes were treated with 50 ng/mL phorbol 12-myristate 13-acetate (PMA; GenScript, USA) for 24 hours prior to co-culture. An indirect co-culture system was established using Transwell inserts with a 0.4 μm pore size polycarbonate membrane (Corning, USA). Briefly, PMA-differentiated THP-1 macrophages were seeded into the lower chambers while BxPC-3 or PANC-1 cells were seeded into the upper chambers. The co-culture system was maintained under standard incubation conditions for 96 hours prior to downstream analyses. Control groups consisted of tumor cells or macrophages cultured alone under identical conditions.

### Small interfering RNA (siRNA) transfection

To transiently knock down LILRB4 expression, THP-1 cells were seeded into 6-well plates and transfected with specific siRNAs using Lipofectamine™ LTX Reagent (Thermo Fisher Scientific, Waltham, MA, USA) following the manufacturer’s instructions. Two independent siRNA sequences targeting human LILRB4, alongside a non-targeting scrambled control, were synthesized by Tsingke Biotechnology (Beijing, China). The specific sequences were as follows: si-R1 (sense: 5’-GGAGUACCGUCUGGAUAAA-3’, antisense: 5’-UUUAUCCAGACGGUA CUCC-3’); si-R2 (sense: 5’-GGAGCCUACAGUAAACCCA-3’, antisense: 5’-UGGGUUUACUG UAGGCUCC-3’); and si-R3 (sense: 5’-GGACUAUGCAGGGAGAUAC-3’, antisense: 5’-GUAUCUCCCUGCAUAGUCC-3’). The specific knockdown efficacy was comprehensively validated by assessing mRNA levels via RT-qPCR at 48 hours post-transfection, and protein expression levels via Western blot at 72 hours post-transfection.

### Western blotting

Proteins were extracted from cells with RIPA buffer supplemented with PMSF and mixed with SDS loading buffer (Beyotime, China) before boiling for 10 min. Then, the proteins were separated by SDS–PAGE and transferred onto a PVDF membrane (BioRad). After blocking the membranes with milk at room temperature for 2 hours, the membranes were incubated with the appropriate antibody at 4 °C overnight. After washing with TBST buffer three times, the membranes were incubated with the corresponding HRP-labeled secondary antibody (Proteintech, China) at room temperature for 2 h, after which they were washed again with TBST three times. Finally, the membranes were exposed and visualized with Chemiluminescence HRP Substrate (Millipore, USA). The primary antibodies used to western blot analysis included: anti-N Cadherin (Abcam, ab98952, 1:1000), anti-E cadherin (Abcam, ab40772, 1:1000), anti-KRT5 (Abcam, ab52635, 1:10000), anti-GATA6 (Cell Signaling, 5851, 1:1,000), anti-P63 (Abcam, ab124762, 1:1000), anti-ERK 1/2 (Abcam, ab184699, 1:10000), anti- phospho- ERK 1/2 (Abcam, ab201015, 1:1000), anti-MEK 1/2 (Cell Signaling, 8727, 1:1000), anti- phospho- MEK 1/2 (Cell Signaling, 9154, 1:1000), anti-beta-Actin (Cell Signaling, 4967, 1:1000).

### PDO establishment and 3D co-culture assay

Patient-derived organoids (PDOs) were established from human pancreatic tumor tissues through enzymatic digestion and subsequent embedding in Matrigel.Dissociated PDO and THP-1 cells were mixed at a 3:1 ratio and co-embedded into Matrigel domes. The 3D direct co-cultures were maintained in organoid medium supplemented with 10 μg/mL of either an isotype control or the LILRB4 monoclonal antibody (hu36B10D1, Nanjing Leads Biolabs Co., Ltd.). Organoid growth and morphological changes were recorded daily over a 7-day period using bright-field microscopy. At the designated endpoint, overall cell viability within the co-culture system was quantified using the CellTiter-Glo® 3D Cell Viability Assay (Promega, USA).

### *In vivo* xenograft models and treatments

All animal procedures were conducted in accordance with the guidelines and approval of the Institutional Animal Care and Use Committee (IACUC) at the First Affiliated Hospital of Nanjing Medical University. Six-week-old NOD-scid IL2rg-/- mice were housed under specific pathogen-free (SPF) conditions. To establish the subcutaneous xenograft models, PANC-1 cells (1×10^6^ cells/mouse) were injected subcutaneously into the flanks of the mice, either alone or co-injected with THP-1 cells (PMA induced M0 status) at a 1:1 ratio in a PBS/ Matrigel mixture. Subsequently, THP-1 cells were pretreated in vitro with either an isotype control or the LILRB4 mAb. Following tumor cell inoculation, mice were randomly allocated into designated treatment groups. Intraperitoneal (i.p.) treatments were initiated on day 6^th^ post-inoculation and administered consistently on days 6^th^, 10^th^, 13^th^, 17^th^, 20^th^, 24^th^, 27^th^, and 31^th^. The treatment arms included: Isotype control, LILRB4 mAb (250 μg/time/mouse), the KRAS G12D inhibitor (HRS-4642, 1.5mg/kg/mouse, Jiangsu Hengrui Pharmaceuticals Co., Ltd.), and a combination of LILRB4 mAb and KRAS G12D inhibitor. At the designated endpoints, the mice were euthanized, and the subcutaneous tumors were carefully excised, photographed, and weighed to evaluate the therapeutic efficacy of the respective treatments.

### Transwell invasion assay

For the invasion assay, Matrigel was diluted in serum-free DMEM (1:3) and coated onto the upper chambers of 24-well Transwell plates. Tumor cells (murine KPC or human PANC-1) were trypsinized, manually counted, and resuspended at a density of 5 × 10^4^ cells/mL in serum-free medium. Subsequently, 100 µL of the respective cell suspension was loaded into the upper chambers. To evaluate paracrine effects, two parallel setups were utilized: (1) for KPC cells, the lower chambers were filled with conditioned medium (CM) derived from FACS-sorted LILRB4^+^ or LILRB4^-^ TAMs; (2) for PANC-1 cells, the lower chambers were filled with CM collected from co-cultures pre-treated with either an isotype control or the LILRB4 monoclonal antibody (hu36B10D1, Nanjing Leads Biolabs Co., Ltd.). Following a 24-hour incubation at 37°C with 5% CO2, the assay was halted, and non-migrated cells in the upper chambers were gently removed using a cotton swab. Cells that had successfully invaded through the Matrigel membrane were fixed with 4% paraformaldehyde, stained with 0.05% crystal violet, and counted under a light microscope. The mean number of invaded cells per membrane was determined by randomly selecting at least five fields and analyzing them using ImageJ software (NIH, USA).

### Mouse model of KRAS inhibitor-resistant pancreatic cancer

By combining high-dose drug with incremental drug concentration, the primary KPC cells were treated with the KRAS inhibitor (HRS-4642) at a concentration of 200 nM. Subsequently, during each subsequent culture, the drug concentration was increased incrementally in a gradient of 10, 50, 100, 200, 400, 800, and 1500 nM until reaching 1500 nM, and then the cells were cultured at this concentration, thus obtaining an in vitro resistant cell model. Approximately 1×10⁶ in vitro resistant cells or the same number of KPC sensitive cells without drug treatment were injected into the orthotopic pancreatic tissue of C57BL/6J mice. After the injection, the formation of pancreatic implanted tumors was monitored every 3 days. After pancreatic orthotopic tumors with a diameter of 1-2 cm were formed in about 2 weeks, the KRAS inhibitor HRS - 4642 (0.03 mg/g) was intraperitoneally injected to further verify the drug resistance. Then, the pancreatic implanted tumors were collected for scRNA sequencing.

### Flow Cytometry Analysis and Fluorescence-Activated Cell Sorting (FACS)

Fresh tumor (T), and normal tissues adjacent to the tumor (NAT) tissues were obtained from human pancreatic cancer patients. Additionally, murine orthotopic pancreatic tumors were established by injecting KPC cells into the pancreata of C57BL/6 mice and were subsequently harvested. Both human and murine tissues were mechanically dissociated and enzymatically digested to generate single-cell suspensions. Following filtration and washing, the cells were resuspended in a flow cytometry staining buffer.

To identify and isolate specific macrophage subpopulations, the cell suspensions were first incubated with fluorescently labeled antibodies targeting macrophage surface markers. Following the necessary washing steps, the 7-AAD viability dye was added to the cell suspensions immediately prior to flow cytometric acquisition. During the gating strategy, dead cells were initially excluded by gating on the living (7-AAD-) cell population. Subsequently, TAMs were identified as CD68^+^ CD11b^+^ cells, whereas murine TAMs were identified as CD11b^+^ cells. Finally, these respective TAM populations were evaluated for LILRB4 expression, and the LILRB4+ and LILRB4- subpopulations were sorted using a flow cytometer for downstream applications.

The antibodies and reagents used for flow cytometry analysis and sorting were included as follows: 7-AAD Viability Staining Solution (Thermo Fisher, 00-6993-50, 5 µL/test). For human TAM sorting, the antibodies included: anti-human CD68 PE (Thermo Fisher, MA5-23572, 5 µL/test), anti-human CD11b FITC (Thermo Fisher, MA1-80592, 5 µL/test), and anti-human LILRB4 (CD85k) APC (Thermo Fisher, 17-5139-42, 5 µL/test). For murine TAM sorting, the antibodies included: anti-mouse CD11b FITC (Thermo Fisher, 11-0112-82, 0.5 µg/test) and anti-mouse LILRB4 (gp49 Receptor) Alexa Fluor® 647 (BioLegend, San Diego, CA, USA; Catalog No. 144905, 0.25 µg/test).

## Data Availability

The newly generated data will be available upon publication. Publicly available spatial transcriptomics data from Kim *et al*^55^. were obtained from the NCBI Gene Expression Omnibus (GEO) under accession GSE235315. Publicly available single-nucleus and spatial profiling data from Hwang *et al*^19^ was obtained from GEO under accession GSE202051. Additional PDAC 10x Visium spatial transcriptomics datasets generated by the Washington University (WUSTL) Human Tumor Atlas Network (HTAN) atlas^20^ were downloaded from the HTAN Data Coordinating Center (DCC) Data Portal.

## Acknowledgements

This work was supported by grants from the National Natural Science Foundation of China (82103235, 82473412), and the Major Basic Research Fund of Jiangsu Province Hospital (TS202407) and the Barnhart Family Distinguished Professorship in Targeted Therapies from the University of Texas MD Anderson Cancer Center. The sponsors or funders were not involved in any parts of this study. The authors thank all contributing physicians, study nurses, and laboratories (Jiangsu Provincial Key Laboratory of Chronic Digestive Diseases) for their support in the study.

