## Supplementary Figures for "Co-evolution of Oncogenic KRAS Signaling and LILRB^high^ Macrophages Drives Pancreatic Cancer Recurrence"

**Supplementary Fig. 1 Related to Figure 1.**

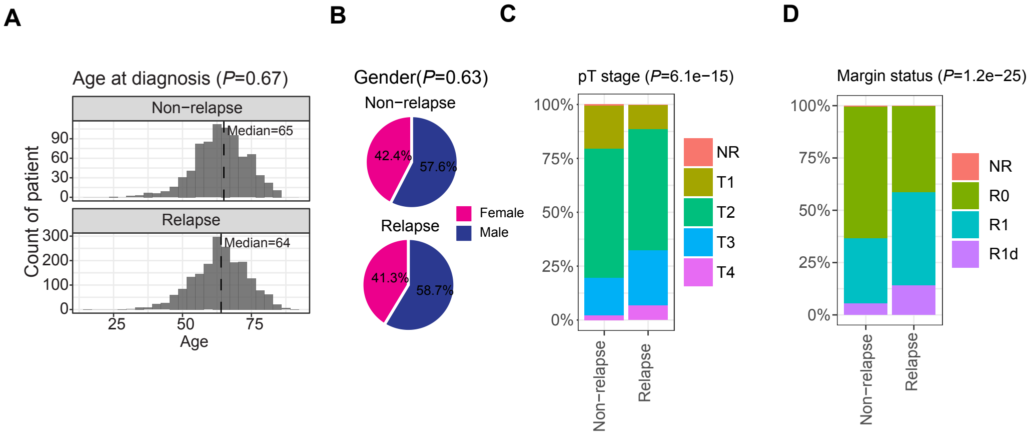


**Supplementary Fig. 1 | Clinical and pathological characteristics associated with recurrence.**

(**A**) Age-at-diagnosis distributions in the non-relapse and relapse cohorts, shown as histograms with median age indicated in each group; *P* value is shown.
(**B**) Sex distribution in the non-relapse and relapse cohorts, shown as pie charts; P value is shown.
(**C-D**) Stacked-bar summaries comparing the non-relapse and relapse cohorts for pathological T stage (C) and margin status (D); *P* values are shown.

PT, primary tumor; RT, recurrent tumor. *P* values are indicated in each panel. Related to **Figure 1**.

#### Supplementary Fig. 2 Related to Figure 2

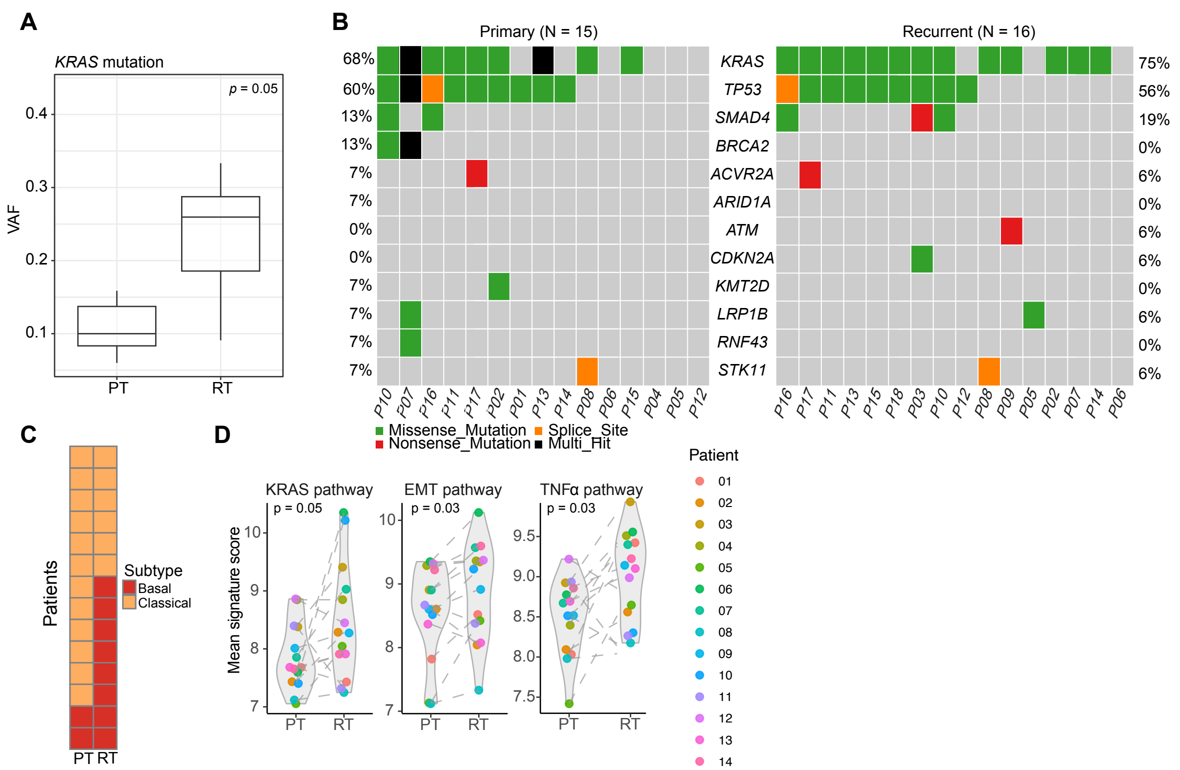


**Supplementary Figure 2 | KRAS mutational and transcriptional changes in paired primary and recurrent tumors**

(**A**) Box plot of *KRAS* variant allele frequency (VAF) in matched primary tumors (PT) and recurrent tumors (RT) from the paired WES cohort; Unpaired t-test *P* value is shown.
(**B**) Oncoprint of recurrently altered PDAC driver genes of the WES data. Columns represent samples and rows represent genes; mutation classes are color coded, and mutation frequencies are shown.
(**C**) Paired subtype quanlitative assignment of tumors (PT versus RT) based on bulk RNA-seq state classification (Classical versus Basal), displayed for individual patients.
(**D**) Paired pathway signature scores for KRAS signaling, EMT, and TNFα programs in PT and RT bulk RNA-seq samples; dots represent individual patients and lines connect matched pairs; P values are shown.

PT, primary tumor; RT, recurrent tumor; WES, whole-exome sequencing; VAF, variant allele frequency; EMT, epithelial–mesenchymal transition. Related to **Figure 2**.

### Supplementary Fig. 3 Related to Figure 3

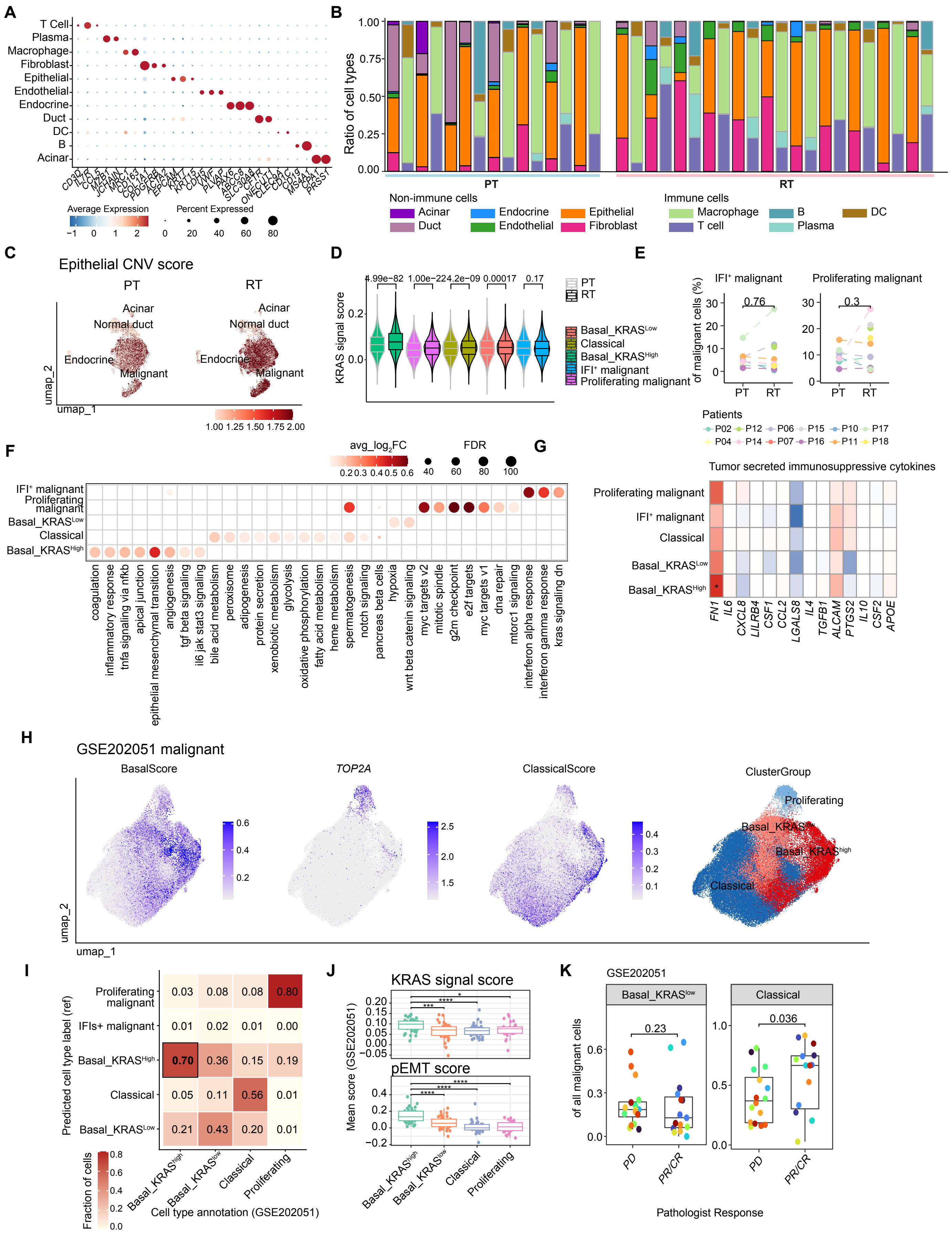

**Supplementary Figure 3 | Single-nuclear characterization of malignant-state composition and external validation of KRAS-activated basal cells.**

(**A**) Dot plot of canonical marker genes used for cell-type annotation across major cell populations in the integrated snRNA-seq dataset. Dot size indicates the fraction of expressing cells and color indicates average expression.
(**B**) Per-sample stacked bar plots showing the relative abundance of major cell types in matched primary tumors (PT) and recurrent tumors (RT).
(**C**) UMAP projections of epithelial cells from PT and RT colored by inferCNV score, used to distinguish malignant cells from non-malignant epithelial populations (including acinar, ductal, endocrine, and normal duct cells).
(**D**) KRAS signaling scores across malignant cell states in PT and RT, including Classical, Basal_KRAS$^{low}$, Basal_KRAS$^{high}$, IFIs$^{+}$ malignant, and Proliferating malignant cells; unpaired t-test *P* values are indicated.
(**E**) Paired quantification of IFIs$^{+}$ malignant and Proliferating malignant cell fractions in PT versus RT across patients; *P* values are shown.
(**F**) Hallmarker functional enrichment summary across malignant states. Dot color indicates average logFC and dot size indicates significance (FDR) for representative pathways/programs.
(**G**) Heatmap of tumor-derived immunosuppressive cytokine expression across malignant states.
(**H**) External validation in public dataset GSE202051: UMAP of malignant cells with feature maps for basal score, *TOP2A*, and classical score, and cluster-group annotation (Basal_KRAS$^{high}$, Basal_KRAS$^{low}$, Classical, and Proliferating).
(**I**) Cross-dataset cell-state concordance between the in-house data derived malignant-state labels and GSE202051 dataset annotations, shown as a fraction matrix.
(**J**) KRAS signaling and pEMT scores across malignant states in GSE202051.
(**K**) Fractions of Basal and Classical malignant cells in GSE202051 stratified by pathologic response (PD versus PR/CR); unpaired t-test *P* values are shown.

PT, primary tumor; RT, recurrent tumor; IFIs, interferon-stimulated genes; pEMT, partial epithelial–mesenchymal transition; FDR, false discovery rate; PD, progressive disease; PR/CR, partial/complete response. Related to **Figure 3**.

#### Supplementary Fig. 4 Related to Figure 4

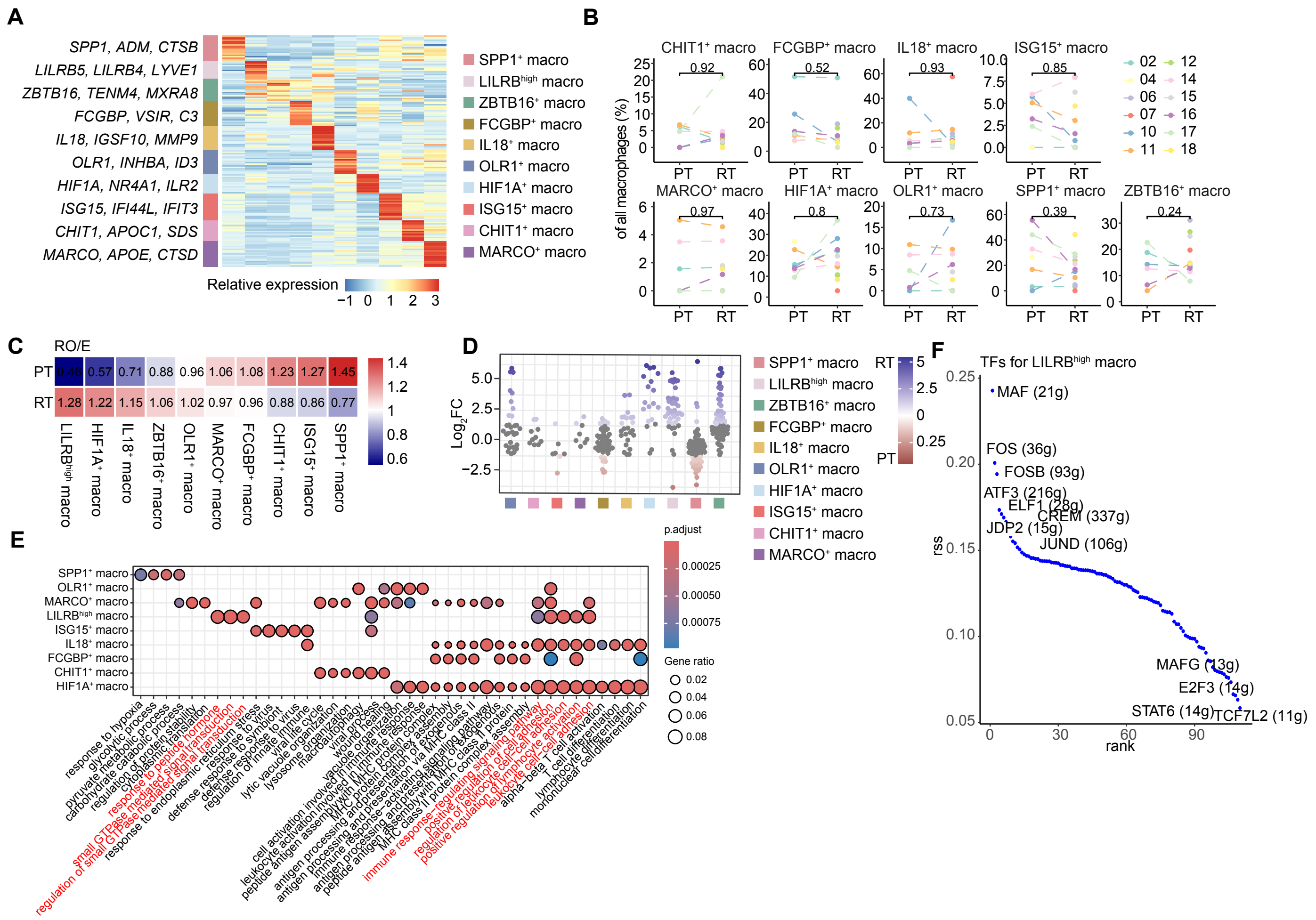


**Supplementary Figure 4 | Macrophage subtype annotation, recurrence-associated abundance shifts, and regulatory features.**

(**A**) Heatmap of representative marker genes across macrophage subtypes identified in the snRNA-seq dataset, showing subtype-defining transcriptional programs (relative expression, scaled by gene).
(**B**) Comparisons of macrophage subtype fractions in primary tumors (PT) versus recurrent tumors (RT) for macrophage states other than LILRBs$^{+}$ macrophages; dots represent samplesand lines connect matched patients; unpaired t-test *P* values are shown.
(**C**) $R_{O/E}$ matrix summarizing relative enrichment of macrophage subtypes in PT and RT (rows, PT and RT; columns, macrophage subtypes), with values shown.
(**D**) Milo differential-abundance analysis on macrophage cells, .
(**E**) Expanded top GO pathway-enrichment dot plot across macrophage subtypes. Dot size indicates gene ratio, and color denotes adjusted *P* value.
(**F**) Ranked transcription factor (TF) regulon activity in LILRBs$^{+}$ macrophages, with selected top or bottom ranked TFs labeled.

PT, primary tumor; RT, recurrent tumor; TF, transcription factor; SCENIC, single-cell regulatory network inference and clustering; RO/E, ratio of observed to expected. Related to **Figure 4**.

#### Supplementary Fig. 5 for Figure 4

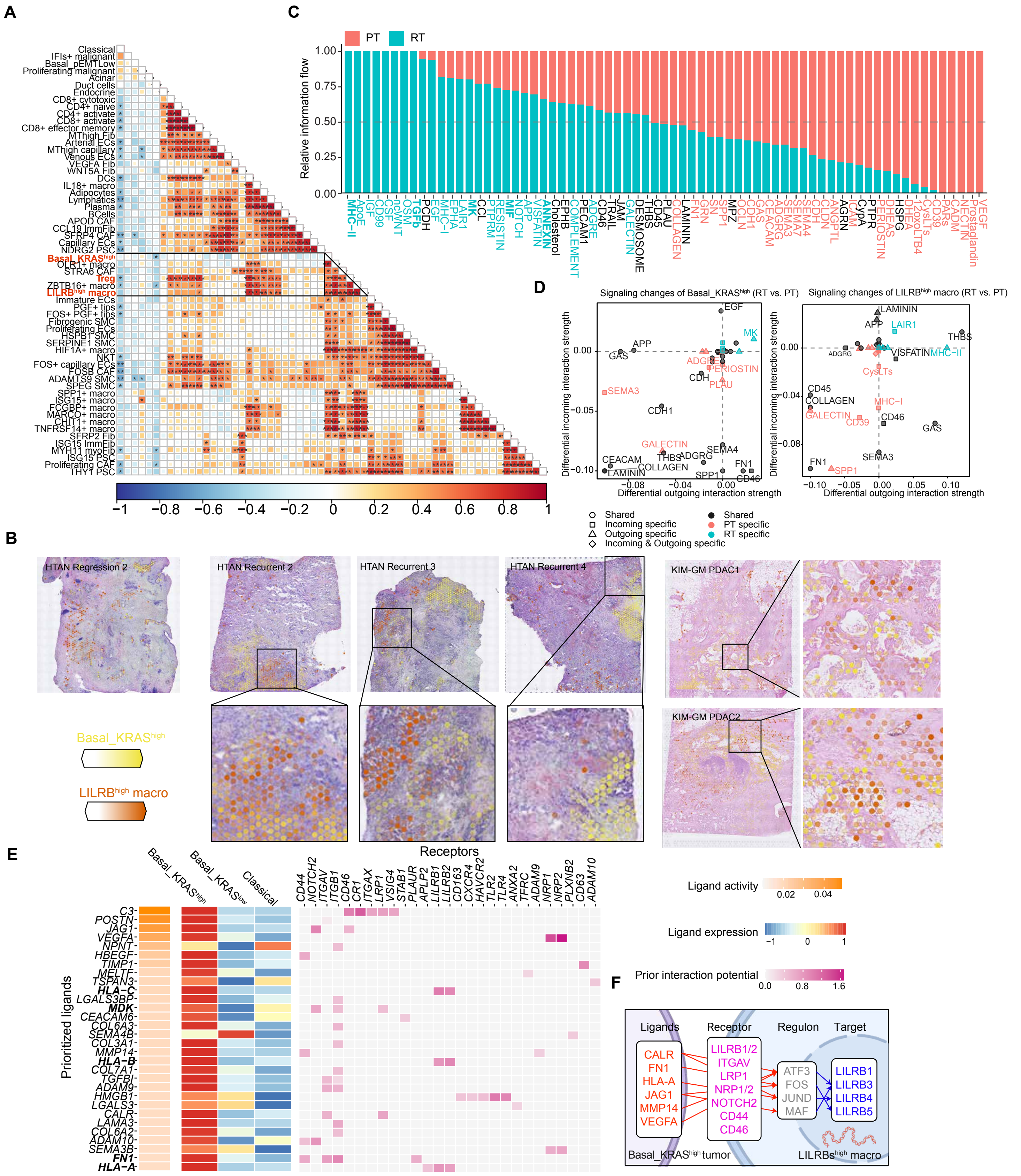

G

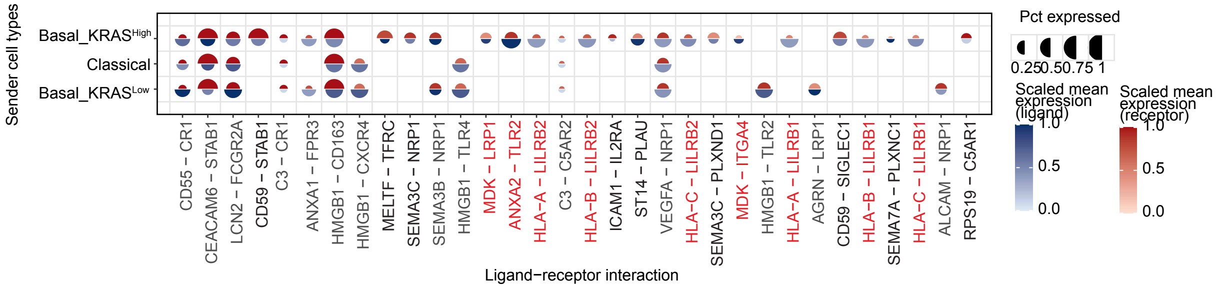


**Supplementary Figure 5 | Cross-compartment coupling of Basal_KRAS$^{high}$ malignant cells and LILRB$^{high}$ macrophages**

(**A**) Pairwise correlation matrix of cell-type abundances across the snRNA-seq cohort, highlighting a co-varying module comprising Basal_KRAS$^{high}$ malignant cells, LILRB$^{high}$ macrophages, and Tregs.

(**B**) Representative spatial maps from external PDAC datasets (HTAN and KIM-GM) showing co-localization of Basal_KRAS$^{high}$ malignant cells (yellow) and LILRB$^{high}$ macrophages (orange), with zoomed-in regions indicated.

(**C**) CellChat information-flow analysis summarizing relative contributions of signaling pathways in RT versus PT; pathways are ordered by overall information flow.

(**D**) Differential outgoing versus incoming interaction strength for Basal_KRAS$^{high}$ malignant cells (left) and LILRB$^{high}$ macrophages (right) comparing RT and PT. Points denote signaling pathways; shared and condition-specific pathways are indicated.

(**E**) NicheNet-based prioritization of candidate tumor-to-macrophage ligands from Basal_KRAS$^{high}$ (and related malignant states), showing ligand activity, ligand expression, and prior interaction potential with macrophage receptors (including LILRB-family and additional receptors).

(**F**) Schematic model of inferred Basal_KRAS$^{high}$ tumor–LILRB$^{high}$ macrophage communication linking prioritized ligands and receptors to macrophage regulons and downstream LILRB-family targets.

(**G**) Dot plot of selected ligand–receptor pairs between malignant states (senders) and macrophage receptors, showing the fraction of expressing cells and scaled mean expression of ligands and receptors.

PT, primary tumor; RT, recurrent tumor; Treg, regulatory T cell. Related to **Figure 4**.

#### Supplementary Fig. 6 Related to Figure 5

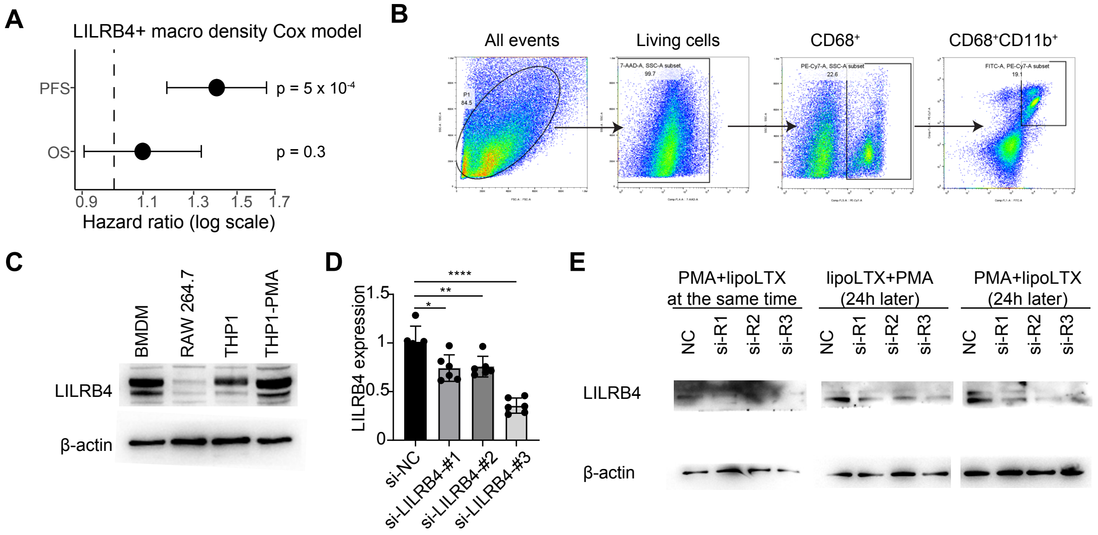


**Supplementary Figure 6 | LILRB4+ Macrophage Density Associates with Survival and LILRB4 Expression Knockdown Is Validated in Macrophage Models**

(**A**) Forest plot showing hazard ratios (HRs) with 95% confidence intervals (CIs) for LILRB4+ macrophage density in relation to progression-free survival (PFS) and overall survival (OS) in the tissue microarray cohort. The dashed vertical line indicates HR = 1. P values from Cox regression are shown. LILRB4+ macrophage density was significantly associated with shorter PFS, but not OS, in this model.
(**B**) Gating strategy for human macrophages.
(**C**) Immunoblot of basal LILRB4 expression in commonly used macrophage/monocyte cell systems (BMDM, RAW264.7, THP-1, and PMA-treated THP-1), with β-actin as a loading control.
(**D**) Analysis of LILRB4 knockdown efficiency in THP-1 cells transfected with control siRNA (si-NC) or independent LILRB4-targeting siRNAs (si-LILRB4-1, si-LILRB4-2, and si-LILRB4-3). Bars show relative expression levels; significance is indicated.
(**E**) Immunoblot validation of LILRB4 knockdown under different THP-1/PMA transfection–differentiation schedules (as indicated), with β-actin as a loading control.

PFS, progression-free survival; OS, overall survival; HR, hazard ratio; CI, confidence interval. BMDM, bone marrow–derived macrophages; PMA, phorbol 12-myristate 13-acetate. Related to **Figure 5**.

**Supplementary Fig. 7 Related to Figure 6**

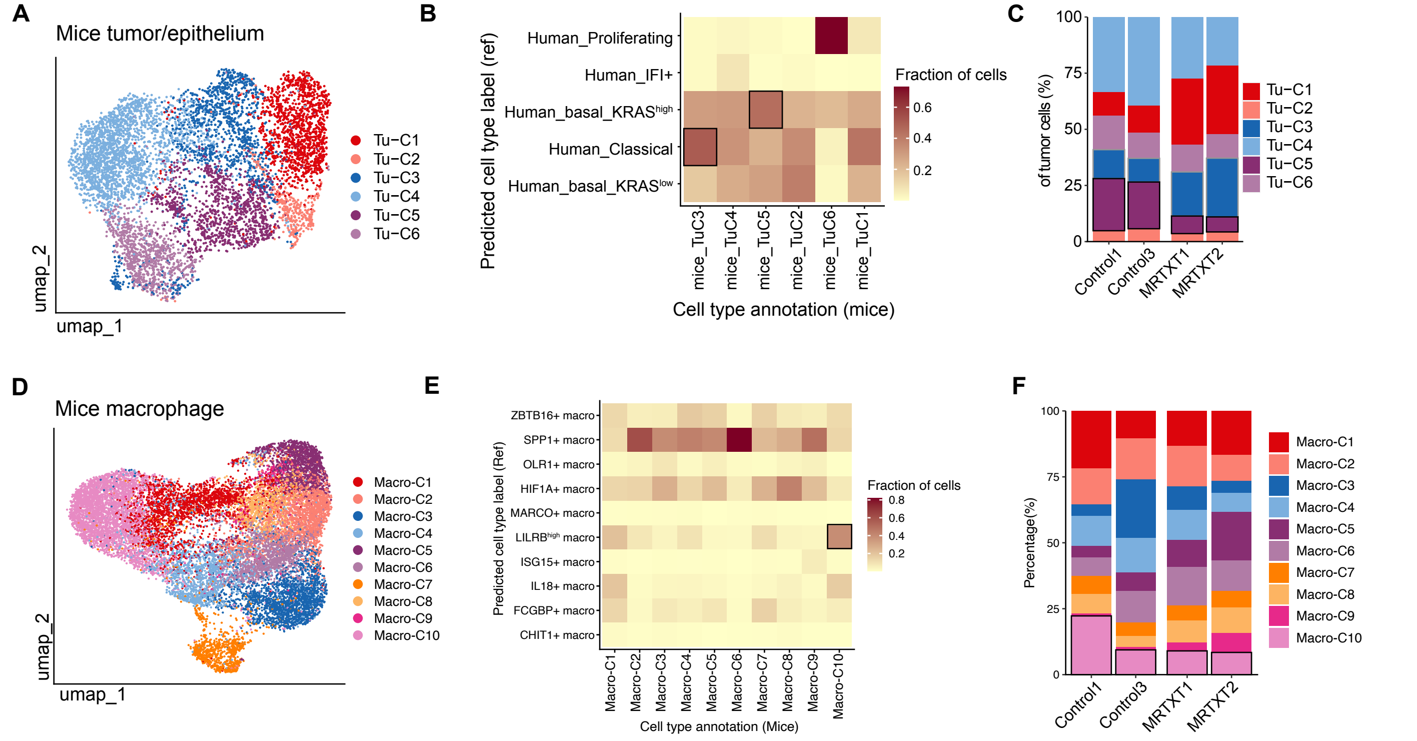


**Supplementary Figure 7 | Cross-species mapping of malignant and macrophage states in the KRAS inhibitor resistance KPC-GEMM PDAC mouse model.**

(**A**) UMAP projection of mouse tumor/epithelial cells, showing six malignant clusters (Tu-C1 to Tu-C6).
(B) Reference-mapping heatmap of mouse tumor clusters to human malignant-cell states . Color indicates the fraction of cells assigned to each reference label; boxed cells mark the dominant match for each state/cluster.
(**C**) Stacked bar plots showing the composition of mouse tumor-cell clusters (Tu-C1 to Tu-C6) across different treatment group samples(Control and treated with MRTXT, as labeled).
(**D**) UMAP projection of mouse macrophages, showing 10 macrophage clusters (Macro-C1 to Macro-C10).
(**E**) Reference-mapping heatmap of mouse macrophage clusters to human macrophage states (including LILRBs$^{+}$ macrophages and other macrophage subtypes). Color indicates the fraction of cells assigned to each reference label; boxed cells indicate dominant cluster-state correspondence.
(**F**) Stacked bar plots showing the composition of mouse macrophage clusters (Macro-C1 to Macro-C10) across individual samples from different treatment groups (Control and MRTXT treatetd).

PDAC, pancreatic ductal adenocarcinoma; UMAP, uniform manifold approximation and projection; IFI, interferon-related; LILRBs+, macrophages with high LILRB-family expression. Related to **Figure 6.**
